# Altered tau in rTg4510 mice after a single interfaced CHIMERA traumatic brain injury

**DOI:** 10.1101/2022.09.01.506145

**Authors:** Wai Hang Cheng, Honor Cheung, Amy Kang, Jianjia Fan, Jennifer Cooper, Mehwish Anwer, Carlos Barron, Anna Wilkinson, Grace Hu, Jefferey Yue, Peter A Cripton, David Vocadlo, Cheryl L Wellington

**Author notes:** **<u>Author Contributions</u>**, Conceptualization: CLW, DV, WHC, Data curation: WHC, HC, AK, JF, JC, CB, AW, GH, JY, Formal analysis: WHC, JF, MA, Funding acquisition: CLW, Investigation; WHC, HC, AK, JF, JC, CB, AW, GH, JY, Methodology: WHC, JF, JC Project administration: WHC Resources; CLW, DV, Software; Supervision: CLW, PAC, DV, Validation; Visualization; n/a, Roles/Writing - original draft: WHC Writing - review & editing: WHC, CLW.

## Abstract

Traumatic brain injury (TBI) in an established risk factor for neurodegenerative disease. In this report, we used the Closed Head Injury Model of Engineered Rotational Acceleration (CHIMERA) to study the effects of a single moderate-severe TBI in rTg4510 mice, a mouse model of tauopathy. Fifteen male rTg4510 mice were impacted at 4.0J at 4-mo of age using interfaced CHIMERA and compared to sham controls. Immediately after injury, moderate-severe TBI induced significant mortality (7/15; 47%), and a prolonged duration of loss of righting reflex. At 2-mo post-injury, surviving mice displayed significant histological evidence of microgliosis (Iba1) and axonal injury (Neurosilver). Western blotting showed that TBI mice had a reduced p-GSK-3β (S9):GSK-3β ratio, suggesting greater tau kinase activity. However, tauopathy in surviving TBI mice showed a divergent response, with 3/8 mice having a very low level of tau protein in brain lysates (i.e. lower than sham), and were thus analyzed separately from the other 5/8 TBI mice that maintained the expected level of total tau. Compared to sham controls, TBI mice with normal tau levels had increased p-tau (PHF1 and AT8), increased autophagolysosome accumulation (p62 and Cathepsin D) and decreased hippocampal size. These findings were not observed in TBI mice with low total tau levels. These observations suggest that TBI leads to chronic white matter injury and altered GSK-3β activity. However, post-injury tauopathy and autophagolysosome accumulation diverged in surviving mice through mechanisms that remain to be defined.

## Introduction

Traumatic brain injury (TBI) is a leading worldwide cause of death and disability. In the USA and Europe, the estimated average annual incidence of hospital-admitted TBI cases is 264 and 262 per 100,000, respectively ^1,2^. A recent study estimated the global incidence of TBI to be 939 per 100,000 ^3^. TBI is clinically classified as mild, moderate, or severe according to the Glasgow Coma Scale, and additional clinical indicators including loss of consciousness and post-traumatic amnesia are valuable for mild TBI (mTBI) that comprise ~90% of reported TBI cases ^4,5^. Repetitive exposure to mTBI is associated with increased risk to develop a neurodegenerative condition called chronic traumatic encephalopathy (CTE) ^6–8^. CTE is a neuropathologically defined disorder, with aggregates of perivascular phosphorylated tau (p-tau) located in the depths of the cortical sulci being its pathognomonic feature ^9^. The p-tau species in CTE include both 3R and 4R tau ^8,10^, indicating both similarities ^10,11^ and differences ^12,13^ with the p-tau species found in Alzheimer’s Disease.

In addition to repetitive mTBI, several reports and case studies show that a single exposure to moderate or severe TBI (msTBI) may also induce p-tau neuropathological changes. A previous report comparing fatal TBI cases to controls (N=5 per age group of <20 yrs, 20-50 yrs, >50 yrs) identified subtle tau immunoreactivity in glial cells during the acute period (<24 hr to 1 month) after msTBI ^14^. A recent study of 39 post-mortem brains of chronic (1-47 years post-injury) survivors of a single msTBI found that, in the young cohort, tau-positive neurofibrillary tangles (NFT) were more frequently observed in TBI cases than in age-matched controls ^15^. Other case studies ^16,17^ examined post-mortem brains from patients who survived decades (24 to 42 years post-injury) after a single msTBI, mostly due to motor vehicle accidents or gunshot, and showed p-tau immunoreactivity and NFT at perivascular sites, superficial and deep cortical layers (sulcal depths), hypothalamus and brainstem. Notably, this study also found evidence of multiple proteinopathies including TDP43. Another recent study ^18^ used positron emission tomography (PET) with the tau tracer flortaucipir to study tau pathology in 21 patients with a remote history of a single msTBI (range 18-51 years post-injury, median 32 years) compared to age-matched controls and found increased flortaucipir-binding at the right lateral occipital cortex in patients with a history of msTBI. In addition, flortaucipir-binding in TBI patients was associated with increased total tau and p-tau concentrations in cerebrospinal fluid (CSF). Together, these studies suggest that a single msTBI may be sufficient to induce long-term tau pathology.

Other case studies have examined post-mortem brains of schizophrenia patients who underwent prefrontal leucotomy ^19,20^. These cases represent a human model of long-term (over 40 years post-injury) single severe traumatic axonal injury. These studies observed perivascular p-tau, NFT and astrocytic tangles only in leucotomized brains, suggesting that p-tau pathologies may be triggered by axonal injury in general and are not necessarily limited to mechanical impact.

Many experimental studies have studied the effects of TBI in various transgenic mouse models that harbour the human *MAPT* gene. Most of these studies focused on relatively “mild” TBI and focus primarily on acute and sub-acute post-injury outcomes. Yoshiyama et al ^21^ induced repetitive closed head impacts (16x, 4 times per day, 1 day per week) to 12-mo Tg T44 mice, which overexpress the shortest human tau isoform T44. At 9-mo post-TBI, only 1 out of 18 injured mice showed extensive NFT development at hippocampus and entorhinal cortex and increased p-tau (PHF1, PHF6, 12E8). Other groups have investigated TBI effects in mice that lack endogenous mouse tau and express all 6 isoforms of human tau (hTau mice), with inconsistent results. Mouzon et al ^22^ induced repetitive closed head impact (5x over 9 days) to 3-mo or 12 to 13-mo hTau mice and observed increased p-tau (RZ3) in the hippocampus at 24 hr post-injury in male but not female mice. Gangoli et al ^23^ performed repetitive un-interfaced CHIMERA mTBI (20×0.24J or 20x 0.13J) to 4-mo hTau mice and observed no p-tau in TBI mice at 1 yr post-injury. Ojo et al ^24^ induced repetitive closed head impacts to 12 wk old hTau mice (24 impacts over 3 months or 32 impacts over 4 months, with an inter-injury time of 72 to 96 hours and observed increased total tau, tau oligomers (TOC-1) and p-tau (pThr231) at 3-mo after the final injury in hTau mice. Other researchers have studied the effects of TBI in mouse models that express mutant forms of tau found in neurodegenerative disease. Tran et al ^25^ performed control cortical impacts (CCI) at 3 severities (mild, mild-moderate, moderate) to 5 to 7-mo 3xTg-AD mice and observed increased tau punctae and increased p-tau (pS199, pT205, pT231, pS396, pS422, AT8, AT100, AT180) in hippocampus, fimbria, and amygdala of the “moderate” injury group from 24 h to 7d post-TBI. Xu et al ^26^ performed weight drop TBI (1x, 4x over 7 days, or 12x over 7 days) in 5 wk-old mice overexpressing P301S tau (1N4R) and observed and injury dosedependent increase of p-tau (pS422) in the retina of TBI mice. Using the same P301S tau model, Edwards III et al ^27^ induced a single msTBI using CCI to 3-mo P301S mice at the right parietal cortex. They observed increased p-tau (AT8) in both ipsilateral and contralateral cortex and hippocampus up to 6-mo post-injury. Interestingly, they also observed tau spreading to the brainstem, and possibly hypothalamus, months after injury. Cheng et al ^28^ induced rmTBI using closed head impact (42 impacts over 7 days) to 3-4 mo Tau58.4 mice that overexpress P301S human tau and did not observe altered tauopathy up to 3-mo postinjury. Bachstetter et al ^29^ induced 1 or 2 closed head impacts 24 hr apart to 2-mo rTg4510 mice that overexpress the P301L human tau (0N4R) and observed increased p-tau (pS396/pS404) by 15 days after the first injury. The discrepancies across these studies likely stem from the many variables in study design, including those focused on the TBI itself (i.e., TBI model, injury severity, injury frequency) as well as in modeling tauopathy (wildtype human tau vs. mutant tau, overexpression vs. endogenous expression level, ratios of different tau isoforms, etc). In general, TBI appears to have a greater probability of inducing tauopathy in mouse models overexpressing mutant forms of tau.

We developed an impact-acceleration model of TBI known as CHIMERA ^30^, which allows unrestrained head motion during head impact. We recently published a modification that allows us to deliver a high impact-energy injury to mice using an interface to distribute impact energy across the skull, thereby protecting from skull fracture ^31,32^. In C57/Bl6 mice, we showed that interfaced TBI at 2.5J resulted in a mortality of <20%, neurological and memory deficits, elevated plasma tau and neurofilament light levels, increased brain cytokine levels and blood brain barrier disruption ^31^. The present study was designed to evaluate the effect of a single msTBI on chronic tauopathy in rTg4510 mice. Here we use interfaced CHIMERA to induce a single msTBI to 4-mo rTg4510 mice, selecting 4.0J to 4.2J impact energy to produce an overall mortality of 46.7%. Animals were aged for 2-mo before brain tissues were harvested for histological and immunoblotting analyses and blood samples were collected for plasma biomarker measurements. We observed that a single msTBI is sufficient to induce long-term white matter injury and prolonged activation of GSK-3β. However, tauopathy and neurodegeneration was exacerbated only in mice with accumulation of p62 and cathepsin D. These findings suggest that disruption of the autophagy-lysosomal pathway may be an important mediator in tau changes after TBI.

## Methods

### Animals

All experiments were approved by the University of British Columbia Committee on Animal Care and are compliant with the Canadian Council of Animal Care (A15-0096). Male rTg4510 mice (Jackson Laboratory #024854) were purchased from the Jackson Laboratory. These mice express a tetracycline-regulatable tetO-MAPT*P301L transgene under the control of the murine prion protein (PrP) promoter, leading to overexpression of human four-repeat (4R0N) mutant P301L tau. These mice also harbour the CaMK2a-tTA transgene, which suppresses tau expression upon exposure to tetracycline or its analogs. Tetracycline was not used in this study. Mice were housed with environmental enrichment on a 12 h/12 h reversed light cycle and received the 2918 Teklad Global 18% Protein rodent diet (Harlan) and autoclaved reverse osmosis water ad libitum.

### Traumatic Brain Injury

At 4-mo (133.4 ±0.5 days) of age, male rTg4510 mice received a single interfaced TBI using the CHIMERA device, as previously described ^31^. Immediately prior to TBI, mice received 0.5 mL NaCl for fluid supplementation and 1 mg/kg meloxicam for analgesia. Anesthesia was induced using 5% isoflurane in 2.5 L/min oxygen and thereafter maintained at 3-4%. Anesthetized mice were restrained by abdominal straps in the supine position on the CHIMERA device, such that their heads were free to move and rested at an angle of approximately 145° relative to the body. A polylactic acid (PLA)-silicone interface fitted to the contour of the mouse skull was placed under the animal’s head to protect from skull fracture and distribute impact energy evenly across the skull. Head acceleration and rotational motion was in the sagittal plane. For sham controls, mice received the fluid supplementation, analgesia, anesthesia, positioning in the CHIMERA device, but no impact. During TBI procedures, the duration of Isoflurane exposure and loss of consciousness were recorded. Impact energies ranged from 2.3J to 5.8J in a pilot experiment designed to define the maximum tolerable impact energy, and an impact energy of 4.0J-4.2J was used for the remainder of this study. Chest compressions with oxygen supplementation was performed on all mice that experienced cardiac/respiratory arrest immediately after the TBI procedure. Most of the animals that did not survive the procedure died within seconds to minutes post-injury. A summary of mortality and causes of death is provided in Supp. Fig. 1A-C.

### Blood Collection and Euthanasia

Longitudinal blood (EDTA-plasma) samples were collected from the saphenous vein 1 week before TBI (defined as baseline), at 6 hr after TBI, and every 2 weeks thereafter, using capillary collection tubes (Fisher). At 2 months post-TBI (6-mo of age), animals were euthanized with 150 mg/kg ketamine (Zoetis) and 20 mg/kg xylazine (Bayer). Cardiac puncture was performed to collect terminal blood samples. Mice were then perfused with 50 mL ice-cold heparinized PBS (5 USP unit/mL). The mouse brain was dissected longitudinally and one hemibrain was frozen for protein homogenization and the other half fixed in 4% paraformaldehyde (PFA) for histology. All blood samples were centrifuged at 1000 *g* for 10 min and the supernatant was stored at −80C as EDTA-plasma.

### Histology, Immunohistochemistry and Immunofluorescence

Hemibrains were fixed in 4% PFA for 2 days and cryoprotected with 30% PBS-sucrose for 3 days, after which 40 μm-thick coronal sections were cut using a cryotome (Leica). Injured axons were stained using the NeuroSilver Staining Kit (FD NeuroTechnologies) following the manufacturer’s instructions. Neurofibrillary tangles were stained using the Gallyas Silver stain protocol, adapted from ^33,34^.

Immunohistochemistry (IHC) for the microglial marker Iba1 was performed as described ^30^. Briefly, sections were quenched with hydrogen peroxide for 10 min, blocked with 5% BSA, and incubated with Iba1 antibody (Wako 019-19741) overnight at 4 °C. Sections were then incubated with biotin-conjugated anti-rabbit secondary antibodies (Biolinx BA1000, 1:1000) for 2 hrs at RT, then with ABC reagent (Vector PK-6100, 1:400) for 1 hr before color development with 3,3’ Diaminobenzidine (DAB) (Sigma D5637). For IHC of tau, total tau was detected with DA9 (1:50,000), and p-tau was detected with AT8 (pS202/T205; Thermo MN1020, 1:500), CP13 (pS202; 1:5000), PHF1 (pS396/S404; 1:5000), and MC1 (pathological conformation; 1:500). DA9, CP13, PHF1 and MC1 were generous gifts from Dr. Peter Davis. Briefly, sections were quenched with hydrogen peroxide for 30 min, blocked with 5% BSA, incubated with primary antibody overnight and then anti-mouse IgG1 (Southern Biotech 1071-08, 1:1000) in 20% Superblock (Fisher PI-37535) for 2 hrs. The sections were then incubated with ABC and developed with DAB as above.

Immunofluorescence of the neuronal marker NeuN was performed by permeabilization of sections with 0.1% Triton X-100 for 30 min, blocking with 5% normal goat serum (NGS), incubation with NeuN antibody (Abcam 104225, 1:1000) overnight at 4 °C, and incubation with anti-rabbit secondary antibody (ThermoFisher A11-012, 1:500) at RT for 2 hrs. For the autophagosome cargo protein p62, sections were permeabilized, blocked, and probed with p62 antibody (NEB D5L7G, 1:800) and streptavidin-594 (Biolegend 405240, 1:500), using the Mouse on Mouse kit (Vector BMK2202) as per the manufacturer’s protocol. The lysosomal proteins cathepsin B and cathepsin D were detected with antibodies NEB DIC7Y (1:1000) and NEB E179 (1:250), respectively. Detection of mouse immunoglobulin G (IgG) was performed using IgG-cy3 antibody (Jackson Immuno Research 715-165-150, 1:50). Co-staining of astrocytes and endothelial cells were performed with antigen retrieval (boiling in sodium citrate buffer in pressure cooker for 5 min) and probing with GFAP-488 (eBioscience 53-9892-80, 1:400) and CD31 (Abcam 28364, 1:200).

### Image Quantification

All coronal brain histology sections were imaged using an Axio Scan.Z1 slide scanner (Zeiss) with a 20x objective. Regions of interest (ROI) include the olfactory bulb, frontal cortex, retrosplenial cortex, hippocampus (dorsal and ventral), amygdala, corpus callosum, optic tract, hypothalamus and ventricles. Quantification of signals from Iba1, NeuroSilver, Gallyas Silver, PHF1, AT8, CP13, MC1, p62, NeuN, and IgG were performed by thresholding and reporting %Area (=Signal Area/ROI Area *100%) ^30,35^. Quantification of GFAP and CD31 signals were performed by reporting mean signal intensity over the ROI and % overlapping area as previously described ^32^. Quantification of CTSB and CTSD signals were performed by reported integrated density over ROI.

### Tissue Homogenization and Western Blot

Proteins from half-brain samples were serially extracted as described ^36^. Briefly, tissues were homogenized in 30 μl/mg of RAB buffer (100 mM MES, 1 mM EDTA, 0.5 mM MgSO4, 750 mM NaCl, 20 mM NaF, 1 mM Na_3_VO_4_, supplemented with protease inhibitor (Roche) and phosphatase inhibitor (Roche). The homogenate was centrifuged at 50,000 *g* for 20 min at 4°C, and the supernatant was collected as the RAB soluble fraction. The pellet was resuspended in 30 μl/mg RIPA buffer (150 mM NaCl, 50 mM Tris, 0.5% deoxycholic acid, 1% Triton X-100, 0.5% SDS, 25 mM EDTA pH 8.0, supplemented with protease inhibitor (Roche) and phosphatase inhibitor (Roche) and centrifuged at 50,000 *g* for 20 min at 4°C, and the supernatant was collected as the RIPA soluble fraction.

Immunoblots were performed by resolving 10 μg of RAB or RIPA fraction through 10% denaturing SDS-PAGE and then transferred to polyvinylidene fluoride (PVDF) membranes (Millipore). Blots were probed by primary and secondary antibodies, developed with Supersignal West Femto (Thermo), and captured by a ChemiDoc MP imaging system (Biorad). The primary antibodies used were: DA9 (1:1,000,000), RZ3 (1:50,000), GluN1 (NEB D65B7, 1:10,000), Synaptophysin (Abcam YE269, 1:50,000), PSD95 (Abcam 18258, 1:100,000), p-GSK-3β (S9) (NEB D85E12, 1:50,000), GSK-3β (NEB 27C10, 1:100,000), GAPDH (Millipore MAB374, 1:100,000). DA9, RZ3, and PHF1 were generous gifts from Dr. Peter Davis. Densitometry was quantified with ImageJ (NIH) using GAPDH for normalization.

### Plasma Total Tau Analysis

Plasma samples from 17 mice were analyzed with the Quanterix^®^ Simoa HD-1^®^ analyzer. Samples were analyzed with the Simoa Tau Advantage Kit (101552) using the manufacturers protocol. Plasma samples were diluted off board at a 50-fold dilution using sample diluent provided. Longitudinal samples were collected from all mice over 6 time points (baseline, 6h, 2 wk, 4 wk, 6wk, 8wk). However, some samples were lost due to machine error. All remaining samples (N=64) were randomized and analyzed on a single plate using the provided 8-point calibrator curve and 2 controls. The curve had an average percent error of 11% and an average recovery of 100%. Both controls were within acceptable ranges as specified by the manufacturer. N=52 samples were run in duplicate, with an average CV of 11%. N=11 samples from 8 mice were run in singlicate due to volume constraints. N=7 samples were over the upper limit of quantification, but within the limits of detection. No samples were below the lower limits of quantification.

### Animal Genotyping

DNA was extracted from ear notches using QIAamp DNA mini kit (Qiagen 51306) following the manufacturer’s instruction. To genotype the Camk2a-tTA transgene, a standard polymerase chain reaction (PCR) assay was performed following the Jax Genotyping Protocol #18439. Specifically, PCR Master Mix (ThermoFisher K0171) and the following cycling conditions were used on a Bio-Rad T100 Thermal Cycler: Initial denaturation at 95°C for 3 min followed by 36 cycles of amplification at 95°C (30 sec), 56°C (30 sec), 72°C (1 min), with a final annealing step at 72°C for 5 min. Samples were kept at 4°C until run on a 1.5% agarose gel (130V). Amplicon size for transgene is ~550 base-pair (bp) whereas the internal positive control gives a band of 324 bp. To genotype the transgene MAPT, probe-specific realtime quantitative PCR (qPCR) was performed using LightCycler^®^ Multiplex DNA Master (Roche07339585001) on a LightCycler^®^ 96 system (Roche) following Jax Genotyping Protocol #22109. qPCR was also used to quantify gene dose of tTa and MAPT in genomic DNA using DNA Green reagents (Roche 06402712001). Primer sequences for tTa are forward: 5’ - GGA CGA GCT CCA CTT AGA CG – 3’, reverse: 5’ - CAA CAT GTC CAG ATC GAA ATC 3’. The same sequences of MAPT and the internal control (APOB) from Jax Protocol #22109 were used to assess gene dose.

### Quantitative Reverse Transcriptase-PCR (qRT-PCR)

RNA was extracted using PureLink^™^ RNA mini-kit (ThermoFisher 12183018A) followed by cDNA synthesis using TaqMan^™^ Reverse Transcription Reagents (ThermoFisher N808-0234) according to the manufacturer’s protocols. Quantitative RT-PCR was performed with DNA Green reagents (Roche 06402712001) on a LightCycler^®^ 96 system (Roche). Each sample was assayed in duplicate and normalized to glyceraldehyde 3-phosphate dehydrogenase (GAPDH). The following primer sequences were used: mouse *Gapdh* forward 5’ - AAG GTC ATC CCA GAG CTG AA - 3’, reverse 5’ - CTG CTT CAC CAC CTT CTT GA - 3’; mouse *Actb* (Actin) forward 5’ - ACG GCC AGG TCA TCA CTA TTG - 3’, reverse 5’ - CAA GAA GGA AGG CTG GAA AAG - 3’; human MAPT (Tau) forward 5’ - CCC AAT CAC TGC CTA TAC CC - 3’, reverse 5’ - CCA CGA GAA TGC GAA GGA - 3’; tTa forward 5’ - GGA CGA GCT CCA CTT AGA CG - 3’, reverse 5’ - CAA CAT GTC CAG ATC GAA ATC - 3’.

### Statistics

In this study, most analyses are comparisons between two groups (sham and TBI). Therefore, depending if data were normally distributed, Student’s t-test or Mann Whitney U test were used for analyses of loss of righting reflex, histology, immunohistochemistry, and Western blot. The exact method used for each plot was stated in figure legend. Longitudinal measurement of plasma total tau was analyzed using log tau level vs log time, using 4 parameter logistic curve model. The effect of impact energy on mortality was modelled using logistic regression model. Correlational analyses of tau vs autophagolysosomal markers were performed using Pearson correlation.

We observed a divergent total tau response in TBI mice, where 3/8 mice had total tau lower than sham controls. Different colours were used to indicate these 3 mice in graphical presentations of TBI group. Specifically, black squares represent the five TBI mice that were found to retain normal levels of total tau (comparable to total tau level in sham animals) upon histological and Western blot analysis, whereas red squares represent the three TBI mice that were found to have a very low level of total tau. Thus, some of the statistical tests we performed analyses twice: with or without the 3 low tau mice. Both p-values were reported in the plots. The upper p-value indicates statistical significance comparing sham vs. all eight surviving TBI mice (i.e. both black and red squares). The lower p-value indicates statistical significance comparing sham vs. TBI mice with expected level of tau (i.e. black squares only). We refer to the upper p-value (comparison between all sham mice and all surviving TBI mice) unless otherwise specified.

## Results

### Mortality and impaired loss of righting reflex after interfaced TBI in 4-mo rTg4510 mice

We first performed a pilot impact energy titration experiment to establish CHIMERA conditions to produce a moderate-severe TBI (msTBI) in 4 month old rTg4510 mice, which overexpress the P301L human 4R0N tau. Interfaced CHIMERA TBI impact energies ranging from 2.3J to 5.8J were tested. Survival was 100% in mice impacted at 2.3-2.9J, 100% at 3.5-3.8J, 53.8% at 4.0J, 50.0% at 4.2J, and 0% at 4.5-4.8LJ (Supp. Fig. 1A). To produce msTBI with an *a priori* defined overall mortality rate of ~50%, we selected an impact energy range of 4.0-4.2J for the full study.

For the full study, seven rTg4510 mice were randomized to the sham group, which received all procedures except for impact, and fifteen rTg4510 mice were randomized to the TBI group and impacted at 4.0 J (N=13) or 4.2J (N=2). Overall, eight mice survived. Six of the thirteen mice that received 4.0J impact and one of the two mice that received 4.2J impact did not survive the procedure. Non-surviving mice either died immediately after TBI or reached humane end point soon after, resulting in an overall mortality rate of 46.7% (7/15) (Fig. 1A, Supp. Fig. 1B, C). The duration of loss of righting reflex (LRR, analogous to loss of consciousness in humans) in all surviving mice is shown in Fig. 1B. On average, sham-injured mice had an LRR duration of 140 sec, whereas TBI mice had a significantly longer LRR duration (2225 sec, p=0.0002).

**Figure 1.**
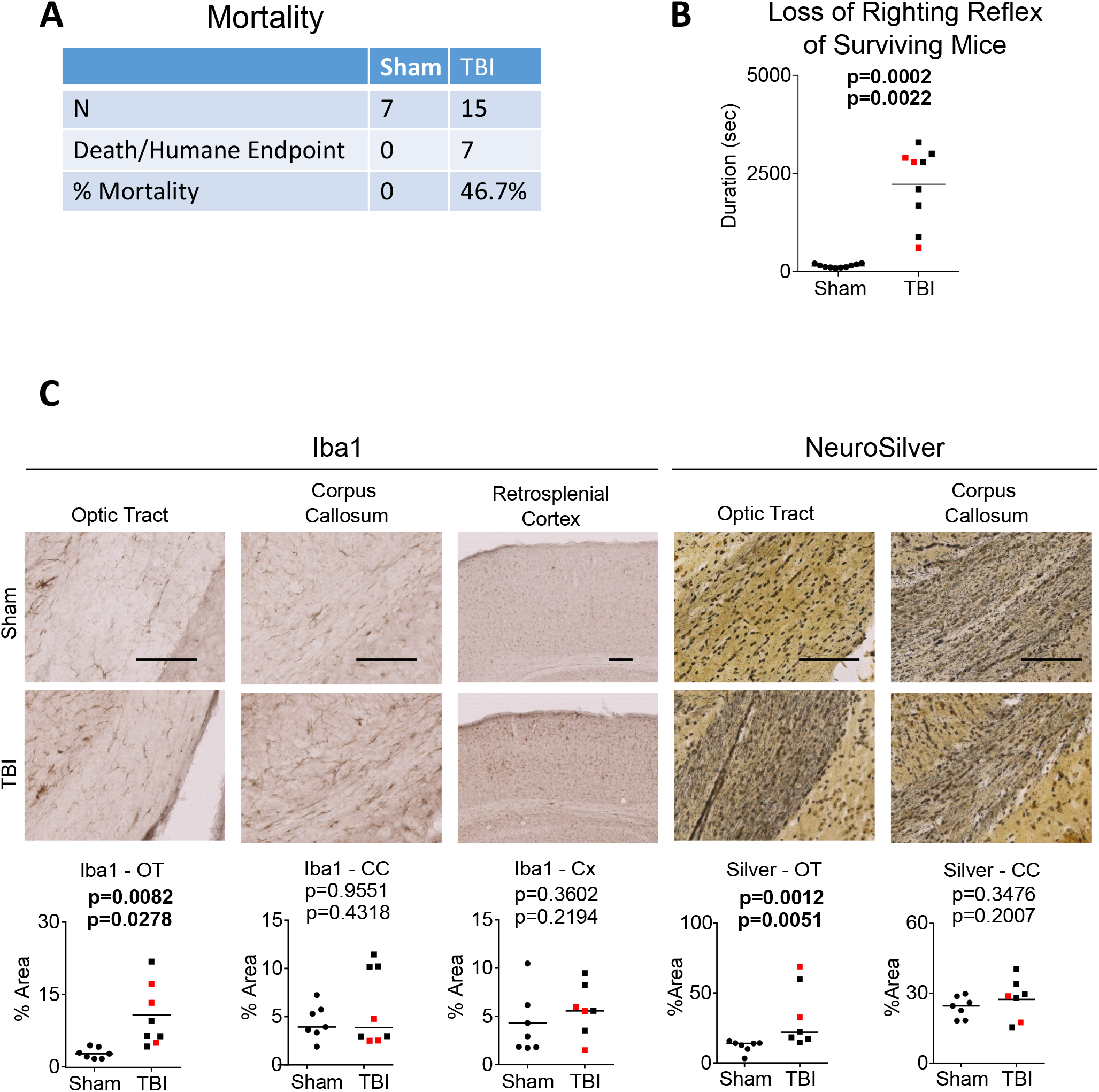
Mortality, loss of righting reflex, axonal injury and white matter microgliosis in rTg4510 after interfaced TBI. (A) Mortality of mice after a single msTBI. (B) Duration of loss of righting reflex of sham and TBI animals. (C) White matter injury in sham and TBI animals as assessed by Iba1 immunohistochemistry in microglia and NeuroSilver staining for degenerative axons. Results are quantified in the graphs below the images. Red squares indicate mice with low tau levels. Scale bar = 100 um. Mann-Whitney U test was used for Iba1-CC and Silver-OT, where horizontal lines indicate group median. T-test was used for all other analyses, where horizontal lines indicate the group mean.

Throughout this report, graphical presentations of rTg4510 TBI data (Figs. 1–7, Supp Fig. 2, 3, 4) include red and black squares, which represent individual animals with divergent responses in many of the tau outcomes. Specifically, black squares represent the five TBI mice that were found to retain normal levels of total tau (comparable to total tau levels in sham animals) upon histological analysis, whereas red squares represent the three TBI mice that were found to have a very low level of total tau upon histological analysis across the panel of antibodies used in this study (Figs. 2A-B). Thus, there are two p-values in graphs comparing sham vs TBI mice. The upper p-value indicates statistical significance comparing sham vs. all eight surviving TBI mice (i.e. both black and red squares). The lower p-value indicates statistical significance comparing sham vs. TBI mice with expected level of tau (i.e. black squares only). We refer to the upper p-value (comparison between all sham mice and all surviving TBI mice) unless otherwise specified.

**Figure 2.**
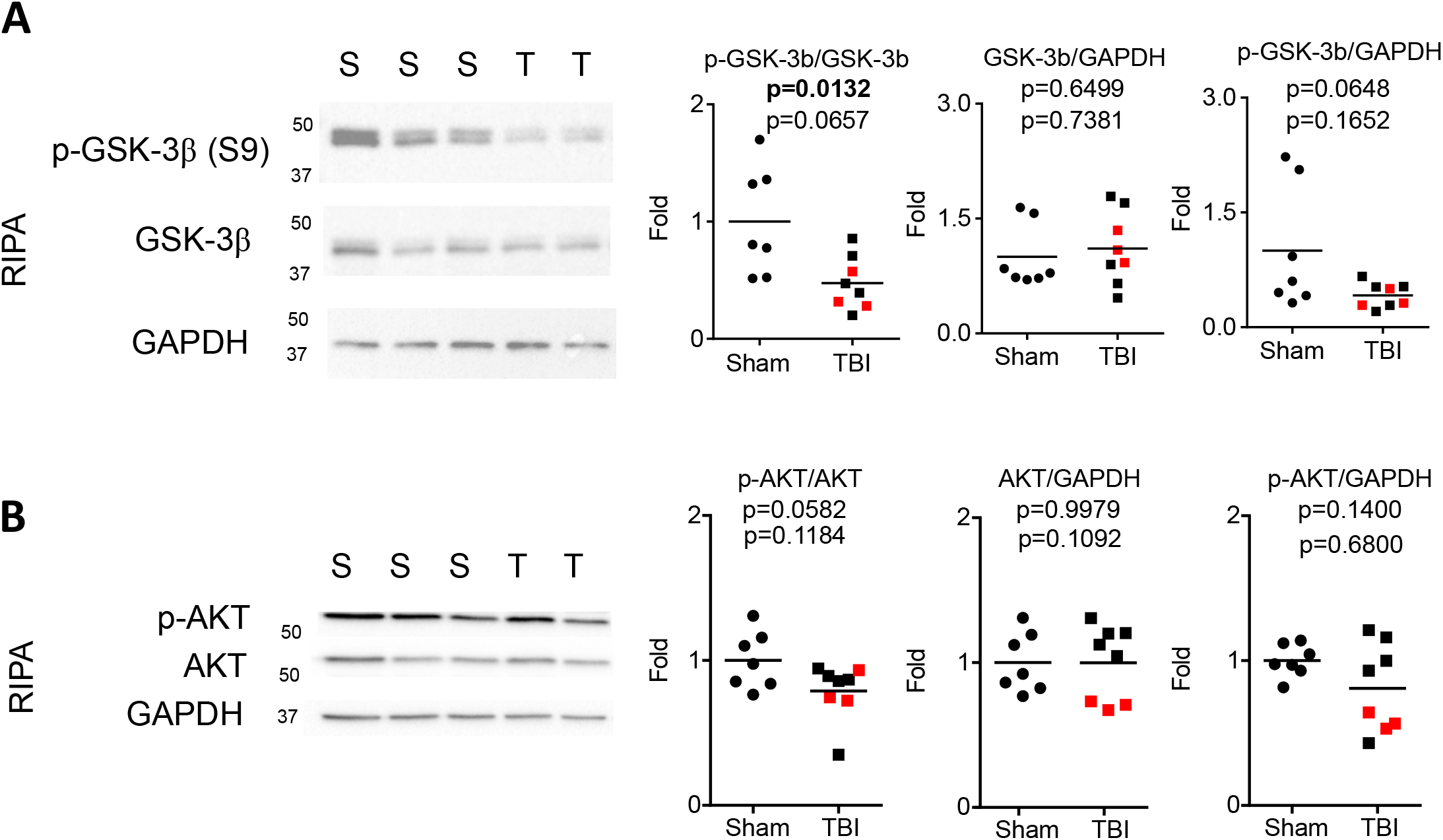
Chronic activation of GSK-3β in rTg4510 after interfaced TBI. Western blotting and quantification of RIPA brain homogenates using antibodies against: (A) GSK-3β and p-GSK-3β (S9), and (B) AKT and p-AKT (S473). Horizontal lines in graphs indicate group mean. Red squares indicate mice with low tau levels. T-test was used for all analyses.

### CHIMERA msTBI induces chronic axonal injury, white matter microgliosis, and activation of GSK-3β in rTg4510 mice

After sham or TBI procedures, the eight surviving rTg4510 mice were aged for 2 months before brains were harvested at 6 months of age. Compared to sham controls, TBI mice had chronic microgliosis at the optic tract as revealed by Iba1 immunohistochemistry (p=0.0082), as well as axonal injury in the optic tract as revealed by silver staining (p=0.0012) (Fig. 1C). These results are consistent with our previous studies demonstrating chronic white matter injury particularly in the optic tract after interfaced CHIMERA impacts in C57Bl/6 mice ^31,35^.

As rTg4510 mice express high levels of 4R P310L human tau, we determined if TBI induced changes in the levels of tau kinases. We first performed Western blotting of RIPA lysates to probe for different forms of GSK-3β, a major Ser/Thr kinase responsible for tau phosphorylation. Although TBI did not affect the level of total GSK-3β (p=0.6499), TBI significantly reduced the level of phosphorylated GSK-3β (p-GSK-3β S9) (p=0.0132), which is an inactive form of GSK-3β (Fig. 2A). This finding suggests that TBI may chronically enhance GSK-3β activity in these animals, potentially increasing p-tau levels after TBI. We then investigated AKT, also known as protein kinase B, which is a Ser/Thr kinase and a major regulator of GSK-3β signaling. When activated, AKT phosphorylates GSK-3β at S9, leading to its inhibition. Western blotting of RIPA lysates revealed no significant change in total AKT levels (p=0.9979) and a strong trend toward reduced p-AKT (S473) levels (p=0.0582) after TBI (Fig. 2B). We conclude that msTBI in rTg4510 mice chronically activates GSK-3β, potentially due to disrupted AKT-GSK 3β signaling.

### Divergent tau changes in rTg4510 mice after CHIMERA msTBI

We next used immunohistochemistry, histochemistry, and Western blotting to evaluate tau burden in all surviving rTg4510 mice (Fig. 3A-B, Fig. 4 A-E). Overall, TBI did not induce significant changes in the level of total tau (DA9, Fig. 3A-B) or neurofibrillary tangles (NFT, by Gallyas Silver stain, Fig. 4A). However, we made the unexpected observation that three of the eight surviving TBI animals showed very low levels of total tau and NFT, which, surprisingly, were lower than tau levels in sham mice in both RAB and RIPA fractions (Fig. 3A-B, Fig. 4 A-E). These “low-tau” mice are highlighted with red squares throughout this report.

**Figure 3.**
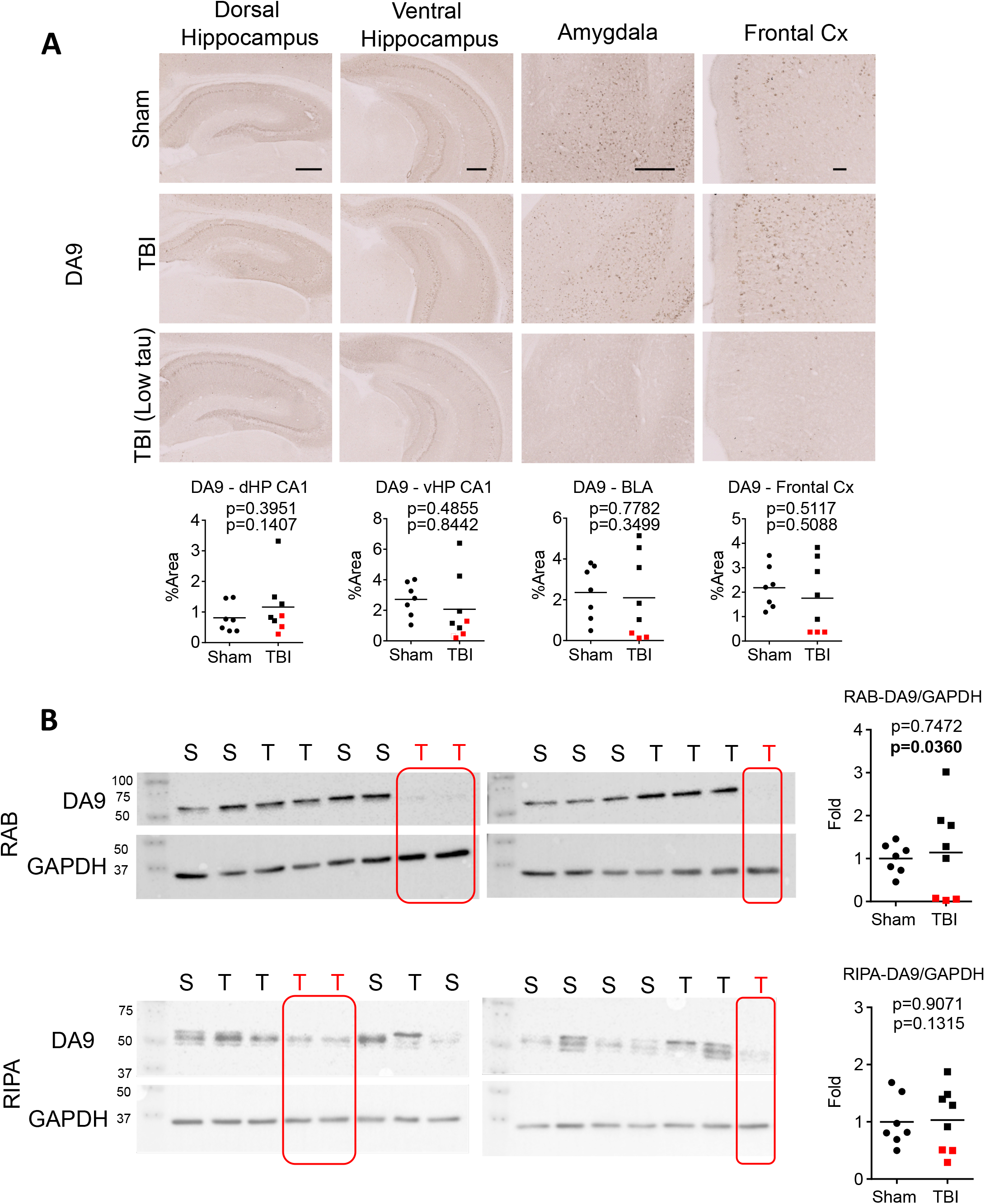
Divergent total tau levels in rTg4510 mice after interfaced TBI. (A) Immunohistochemistry and quantification of total tau performed using DA9. (B) Western blotting and quantification of total tau in RAB and RIPA fractions of brain homogenates using DA9. All sham and TBI samples are shown. The three TBI samples that consistently showed low total tau levels are highlighted in red. Horizontal lines in graphs indicate group mean. Red squares indicate mice with low tau levels. Scale bar = 100 um. T-tests were used for all analyses.

**Figure 4.**
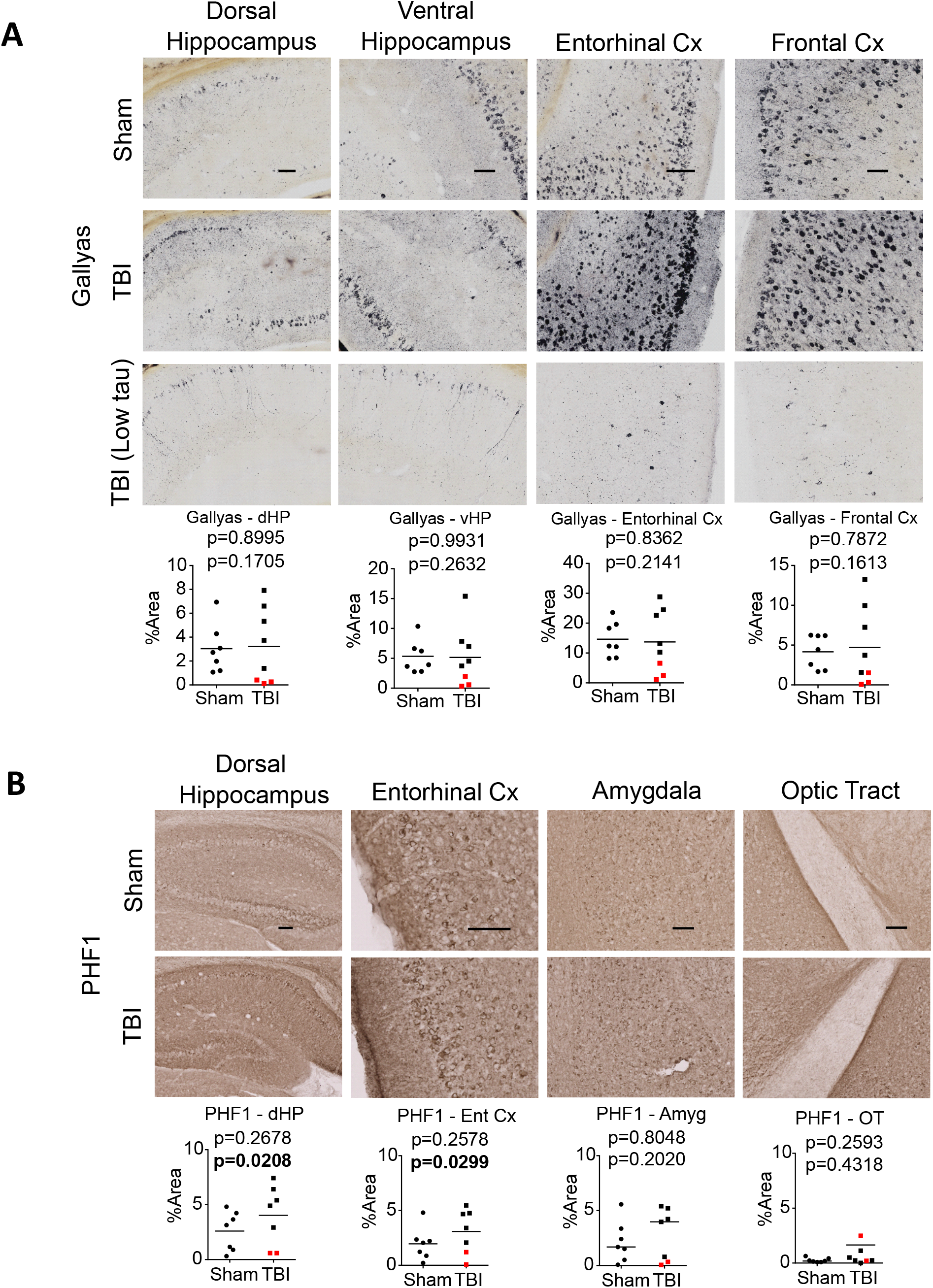
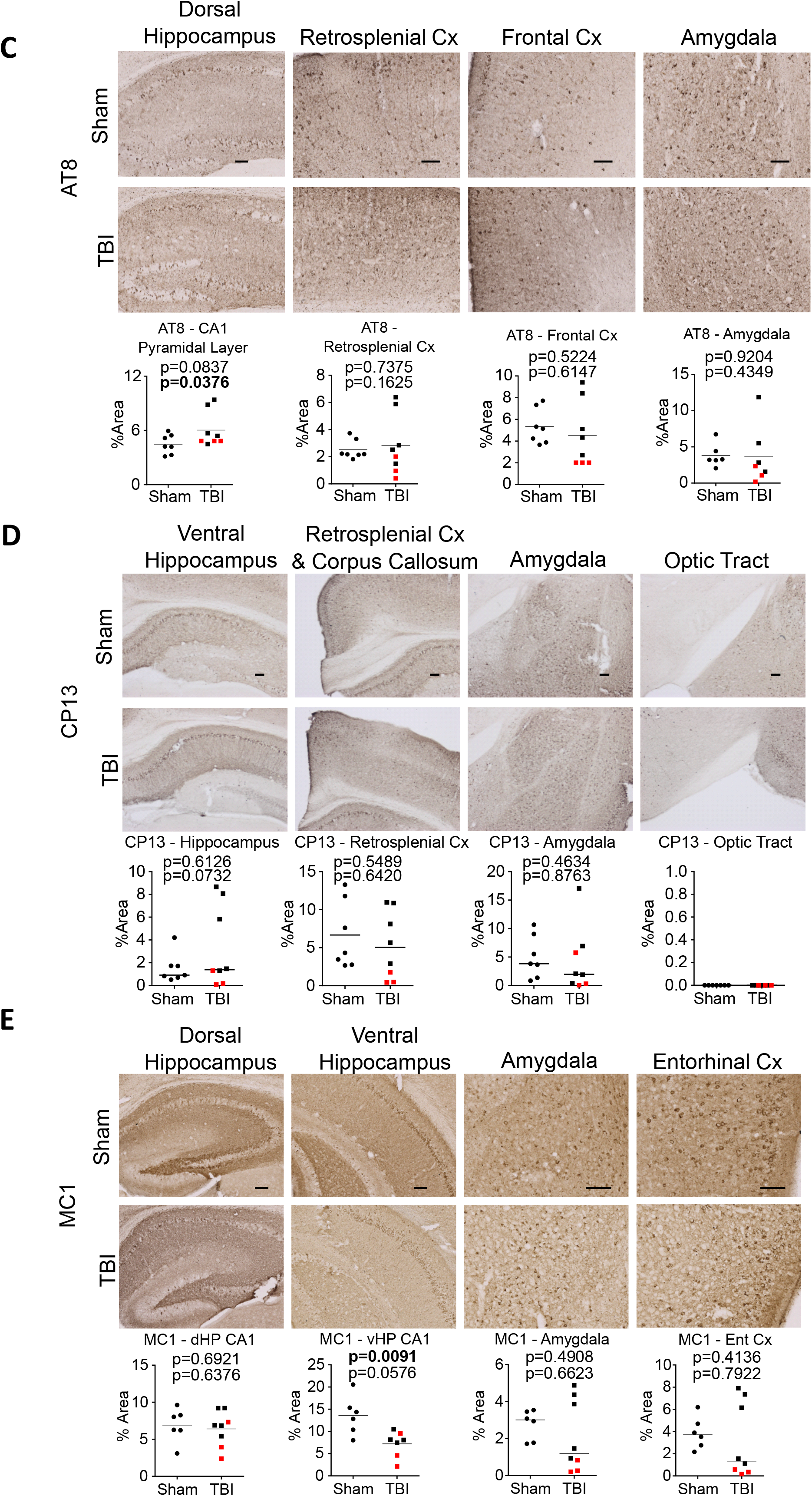
p-tau immunohistochemistry in rTg4510 mice after interfaced TBI. (A) Gallyas Silver staining as a measure of neurofibrillary tangles. (B-E) Immunohistochemistry of p-tau performed using antibodies that recognize different p-tau epitopes: PHF1 (B), AT8 (C), CP13 (D), MC1 (E). Results are quantified in the graphs below the images. Red squares indicate mice with low tau levels. Scale bar = 100 mm. Mann-Whitney U tests were used for PHF1-OT, CP13-HP, CP13-Amyg, MC1-Amyg, MC1-EC, where horizontal lines in graphs indicate group median. T-tests were used for all other analyses, where horizontal lines in graphs indicate group mean.

To further investigate this unexpected finding, we first re-confirmed rTg4510 genotypes (Supp. Fig. 2A). All rTg4510 mice (with normal or low total tau levels) had the correct genotype and were also confirmed to contain the CamK2-tTA (tetracycline-transactivator) transgene. As Gamache et al. ^37^ reported that rTg4510 mice harbour multiple copies of the two transgenes MAPT (for P301L 4R0N tau) and tTA, we next performed quantitative polymerase chain reaction (qPCR) to assess transgene dosage using DNA and transgenic expression using mRNA (Supp. Fig. 2B-D). No significant differences were observed in transgene dosage or transgene expression. We also confirmed there were no tetracycline or related antibiotics used in our animal facility. Having thus ruled out genetic and expression level differences in mice with normal (black squares) and low (red squares) tau levels, we conclude that low tau levels in some mice after TBI is likely due to post-translational mechanisms.

We next performed IHC using antibodies that recognize different phosphorylated tau epitopes: PHF1 (pS396+pS404), AT8 (pS202+pT205), CP13 (pS202), and MC1 (pathological conformation) (Fig. 4B-E). When we compared sham vs all TBI animals, we did not observe a significant increase in any p-tau epitope in any brain region examined (dorsal hippocampus, ventral hippocampus, entorhinal cortex, frontal cortex, amygdala, optic tract, hypothalamus). A significant decrease in MC1 immunoreactivity was found in the CA1 region of the ventral hippocampus of TBI mice (Fig. 4E, p=0.0091), however, this observation was not replicated in any other brain region. Interestingly, when we excluded the low tau mice (red squares) we observed that TBI significantly increased PHF1 immunoreactivity at the dorsal hippocampus and entorhinal cortex (Fig. 4B, p=0.0208 and p=0.0299, respectively). Similarly, after exclusion of low tau mice, TBI also significantly increased AT8 immunoreactivity at the dorsal hippocampus (Fig. 4C, p=0.0376). No differences were found for CP13 after exclusion of low tau mice (Fig. 4D). At the hypothalamus, TBI significantly increased MC1 immunoreactivity after exclusion of low tau mice (Supp Fig. 6 p=0.0173). These findings suggest that while all TBI mice showed comparable injury responses in terms of LRR duration, white matter injury, and GSK-3β activity, the tau outcome was not consistent. Most TBI animals developed more p-tau, but a few had less total and phosphorylated tau across all antibodies used in this study.

### Increased p-tau associates with p62 accumulation, increased cathepsin D levels and decreased hippocampal size

We next tested for changed in the tau degradation pathway to begin to understand the divergent tau response. Using IHC, we probed with an antibody against p62, a protein residing within late endosomes that functions as an autophagosome cargo adaptor protein. Although p62 levels were not significantly different in dorsal hippocampus, ventral hippocampus, frontal cortex and amygdala when shams were compared to the total TBI group (Fig. 5), it is clear p62 levels are lowest in low tau mice in all regions examined. After excluding low tau mice, post-TBI p62 levels were significantly increased in the dorsal hippocampus (p=0.0177) and trends toward increased p62 levels were observed in the ventral hippocampus (p=0.2020), frontal cortex (p=0.2677) and amygdala (p=0.1490). This observation raises the hypothesis that impaired autophagy degradation may lead to increased p-tau levels in mice that retained normal tau levels after TBI. Notably, if we only compared mice with reduced total tau (red square) to sham injured mice, they showed significantly less p62 accumulation in several regions (frontal cortex, amygdala, ventral hippocampus: p=0.0167; dorsal hippocampus p=0.0677).

**Figure 5.**
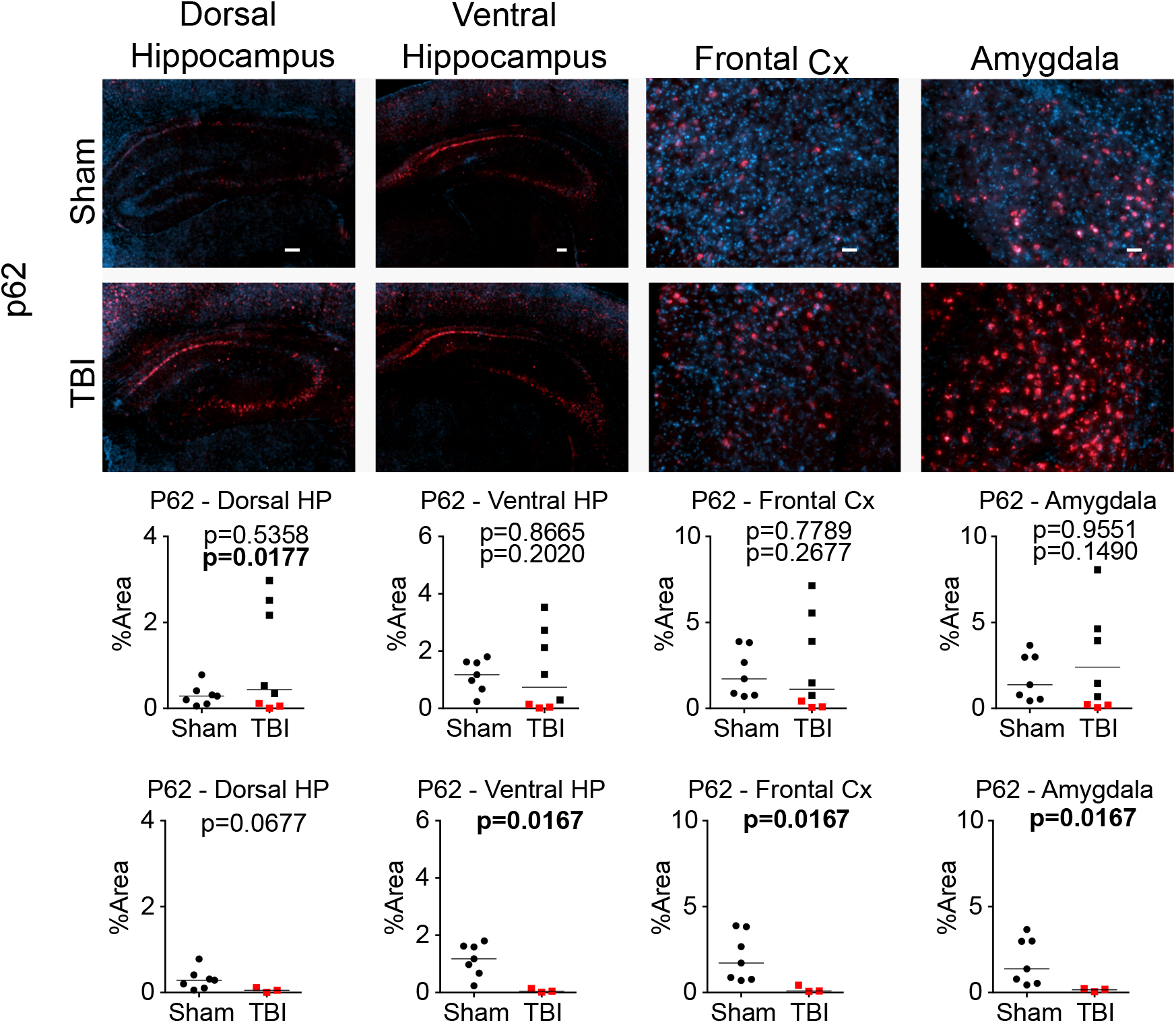
p62 accumulates after interfaced TBI only in rTg4510 mice with normal tau levels. Immunohistochemistry of p62 to stain autphagosomes. Results are quantified in the graphs below the images. Red squares indicate mice with low tau levels. Scale bar = 100 um. Mann-Whitney U test were used for all analyses. Horizontal lines in graphs indicate group median.

We next used IHC to probe cathepsin D and cathepsin B, the two major lysosomal proteases that are involved in proteolysis of tau and other neurodegenerative proteins ^38,39^. We found no significant change of either cathepsin D or cathepsin B immunoreactivity in any area including dorsal and ventral hippocampus and amygdala (Fig. 6A), and frontal cortex (not shown). We also performed Western Blotting for cathepsin D using whole brain RIPA homogenates (Fig. 6B). Although a comparison of sham vs all TBI animals showed no significant change, exclusion of low tau mice revealed increased cathepsin D (p=0.0460) specifically in mice with normal tau levels. A similar trend was found when we analyzed only the mature form of cathepsin D (p=0.0636), although this did not reach statistical significance. This observation suggests that impairment of the autophagy-lysosome pathway in mice that retain normal tau levels is associated with increased p-tau after TBI.

**Figure 6.**
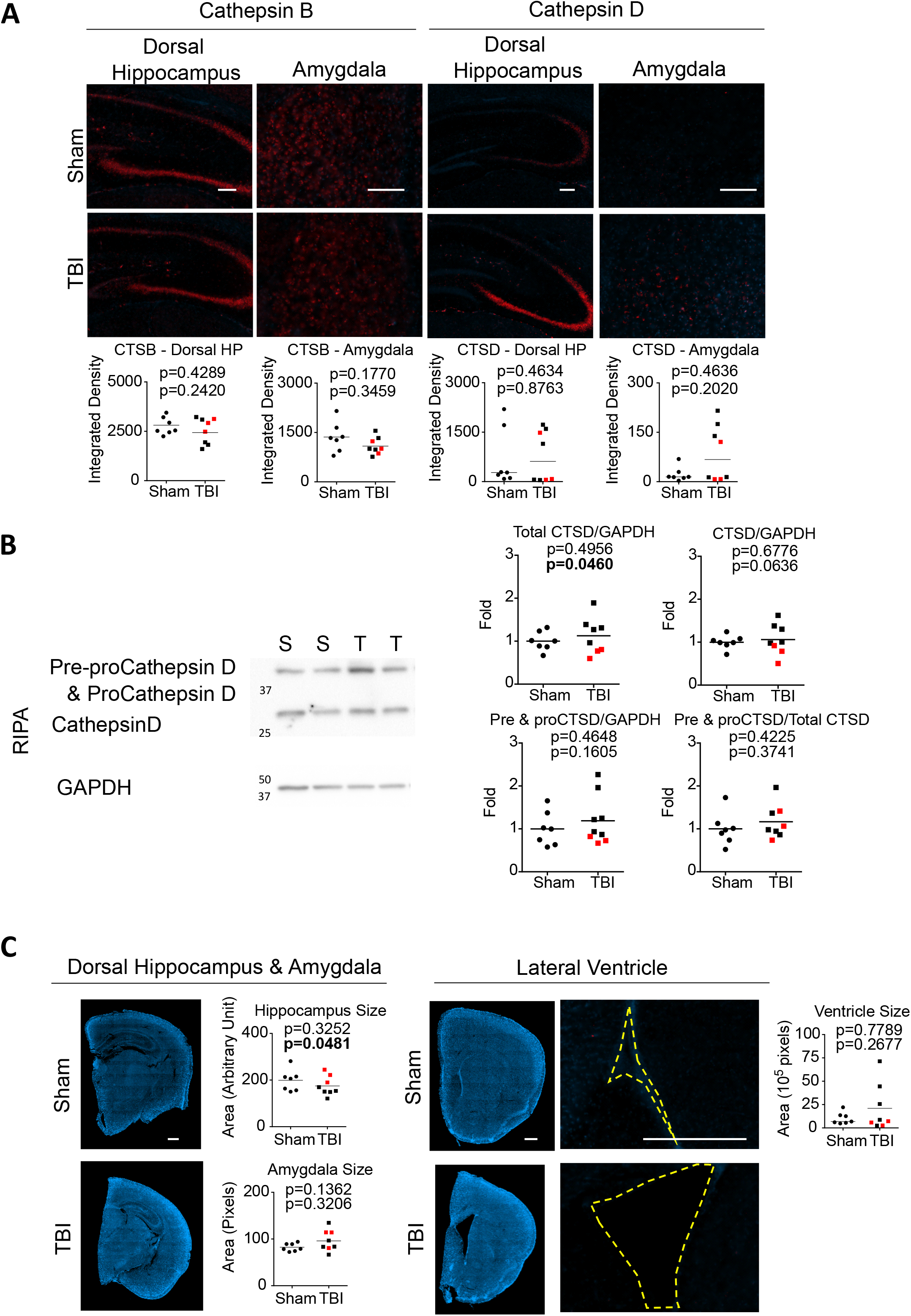
Increased cathepsin D levels and decreased hippocampal size are observed after interfaced TBI only in rTg4510 mice with normal tau levels. (A) Immunohistochemistry of cathepsin B and cathepsin D to stain lysosomes. (B) Western blotting of Pre&pro-cathepsin D and mature cathepsin D performed in RIPA brain lysates. (C) Size comparison of hippocampus, amygdala, and lateral ventricle in sham and TBI mice. Results are quantified in the graphs below the images. Red squares indicate mice with low tau levels. Scale bar = 100 um in (A) and 500 um in (C). Mann-Whitney U tests were used for analyses of CTSD-dHP, CTSD-Amyg, Ventricle size, where horizontal lines in graphs indicate group median. T-tests were used for all other analyses, where horizontal lines in graphs indicate group mean.

We further performed correlational analyses between brain tau burden (p-tau, total tau, or Gallyas silver from cortical IHC or brain homogenate WB) versus markers of autophagolysosomal accumulation (p62 cortical IHC, CTSD/GAPDH WB) or tau kinase activity (p-GSK-3β/GSK-3β WB) (Suppl. Fig. 5). There is a strong and significant positive correlation between cortical level of P62 and tau (R^2^= 0.3139, 0.7093, 0.7140, 0.4033, 0.6668, 0.6403; p= 0.0372, <0.0001, 0.0001, 0.0110, 0.0002, 0.0003, respectively for PHF1, AT8, MC1, CP13, Gallyas, DA9 cortical IHC). A similar positive correlation of p62 and p-tau has been observed in NFT of AD patients ^40^. There is also a significant positive correlation between brain cathepsin D levels and tau (R^2^= 0.3322; p= 0.0310 for PHF1 cortical IHC). However, there is no significant correlation between the ratio of p-GSK-3β/GSK-3β versus tau.

We also analysed the size of various brain regions and ventricles in sham vs. TBI animals (Fig. 6C). The size of the dorsal hippocampus and amygdala were measured in the coronal plane (approximately −2 mm posterior to bregma). The area of the lateral ventricle was measured at the coronal plane of frontal cortex (approximately 1.2 mm anterior to bregma). There was no significant difference in size of these regions when we compared sham vs all TBI animals. However, when we excluded low tau mice, we observed significantly reduced hippocampal size (p=0.0480) specifically in TBI mice with normal tau levels.

### Levels of protein markers of neurons, astrocytes, endothelial cells, autophagy initiation, or synapses are not altered after msTBI

IHC analysis using NeuN, GFAP, CD31, and IgG antibodies to stain for neurons, astrocytes, endothelial cells, and extravasated IgG, respectively, revealed no significant change when comparing sham vs. TBI brains across any region analysed (Supp. Fig. 3). Western blotting also revealed no change in protein levels for autophagy (Beclin1, LC3II/LC3I) or synaptic (synaptophysin, PSD95, and GluN1) markers (Supp. Fig. 4).

### MsTBI may accelerate increase of plasma total tau in rTg4510 mice

We collected longitudinal plasma samples from sham and TBI mice starting from baseline (2 weeks before TBI), at 6 hr post-TBI, and every 2 weeks thereafter. Plasma total tau (t-tau) was assayed using a human t-tau SIMOA assay. Despite loss of some samples due to machine error, we observed a trend toward higher plasma t-tau in TBI mice, including both low tau and normal tau mice, from 2 - 8 weeks post-TBI (Fig. 7A). Four parameter logistic regression was performed on these data (Fig. 7B). The best fitted models of sham and TBI groups did not different greatly in terms of the top plateau (10^2.637^ = 434 pg/ml vs 10^2.733^ = 541 pg/ml). However, we noted that the sham group has a point of inflection at 17 days post-sham procedure (10^1.497^ = 31 days post-baseline), whereas the TBI group has a point of inflection within 1 day post-TBI (10^1.170^ = 14.8 days post-baseline). This finding suggests that msTBI may accelerate the increase of plasma total tau. However, as low tau and normal tau mice performed similarly in the plasma t-tau assay, peripheral sources of t-tau, likely released during neck flexion during the CHIMERA TBI procedure, may confound these results.

**Figure 7.**
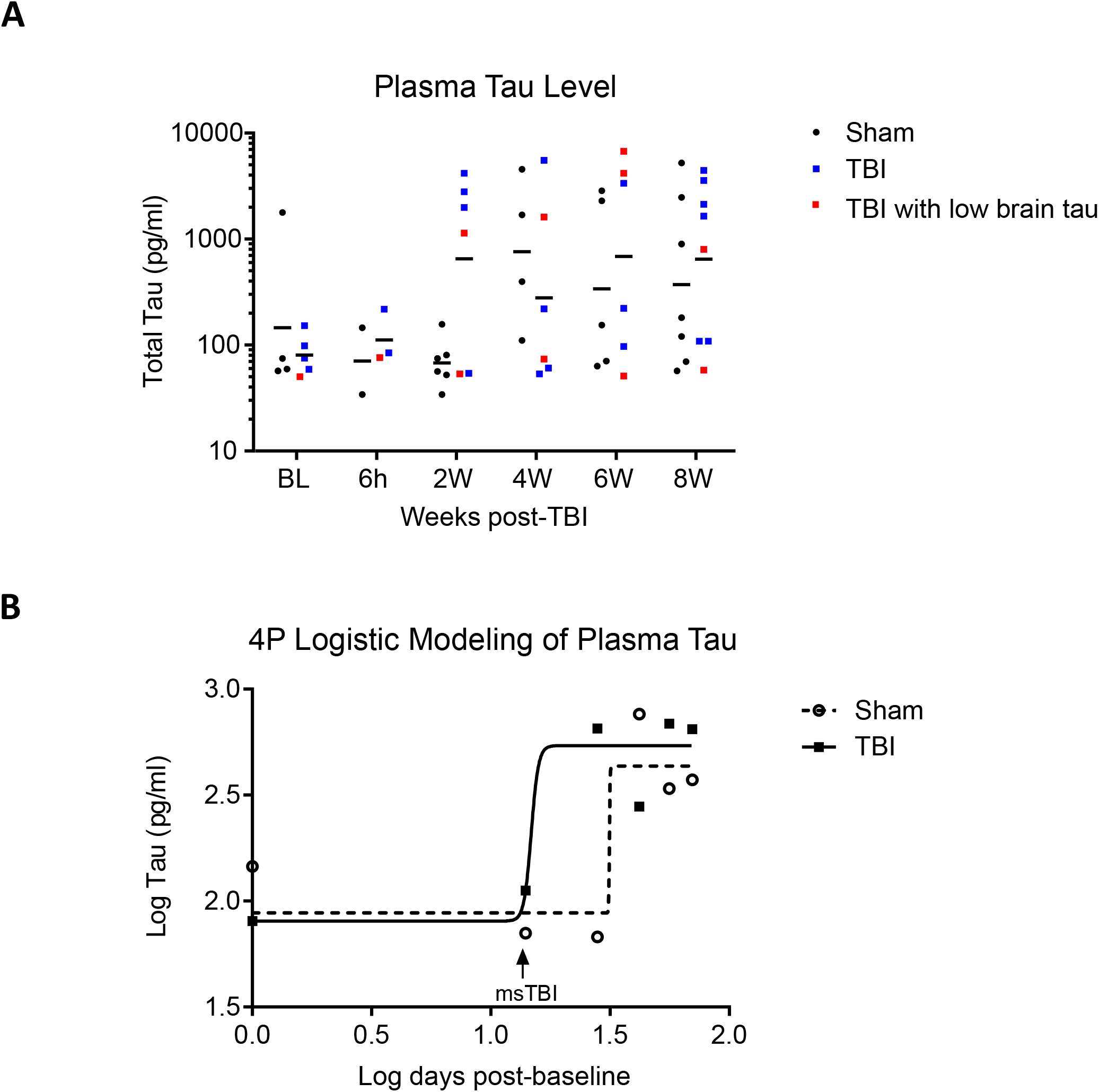
Longitudinal measurement of plasma total tau in rTg4510 mice. (A) Baseline (BL) plasma was collected from rTg4510 mice 2 weeks before TBI. Post-TBI plasma was collected at 6hr post-procedure, and every 2 weeks thereafter. Plasma total tau was quantified using SIMOA. Horizontal line indicates group mean. Red squares indicate mice with low tau levels. (B) Log plasma total tau vs log time was modelled using 4 parameter logistic (4PL) curves. Symbols overlaying the curves represent mean of log values. The sham group was best fitted with the equation y = 1.944 + 0.6926/(1+10 ^326.9 (1.497 - x)^), R^2^ = 0.2389. The TBI group was best fitted with the equation y = 1.905 + 0.8281/(1+10 ^28.91 (1.170 - x)^), R^2^ = 0.1836.

## Discussion

This study was designed to investigate outcomes of a single msTBI induced by interfaced CHIMERA in rTg4510 mice. At an impact energy of 4.0-4.2J, the overall mortality rate in TBI mice in this study was 46.7%, which is aligned with clinical msTBI mortality rates reported at 21% to 43-46% or above ^41–43^. In rTg4510 mice, a single msTBI induced significant acute injury response, namely LRR, as well as chronic white matter injury detectable at 2-mo post-TBI, similar to our previous findings ^31,35^.

A major objective of this study was to determine if msTBI delivered via interfaced CHIMERA exacerbates tauopathy in the rTg4510 model. Clinically, evidence of p-tau accumulation has been reported in case studies of rmTBI, msTBI or axonal injury ^7,8,16,19^. Human tau contains more than 40 phosphorylation sites that are associated with physiological or pathological significance ^44^. Many of these phosphorylation sites have been found in CTE cases ^45^. The phosphorylation and dephosphorylation of tau is regulated by tau kinases and phosphatases. Glycogen synthase kinase (GSK) 3β, which phosphorylates tau at multiple sites ^44,46^, is one of the most well-studied tau kinases. The active form of GSK-3β has been found in NFT of AD patients ^47^, and its inhibition is an active area of therapeutic research for neurodegenerative diseases ^48^. In rodent models of TBI ^49–51^, increased GSK-3β phosphorylation at serine 9 (which inactivates GSK-3β activity ^52^), has been observed during the acute post-injury phase and may be important in neuronal survival in the early period. However, the role of GSK-3β in the chronic post-injury phase is less understood. Interestingly, our study suggests that msTBI chronically disrupts AKT-GSK-3β signaling, leading to increased GSK-3β activity. This finding contrasts with acute rodent TBI studies (hours to days post-TBI) ^49,50,53^. Our observation highlights the potential benefits of inhibiting or competing against GSK-3β activity after TBI, in agreement with reports from other groups using lithium treatment and O-GlcNAcylation^54–59^.

The most interesting and unexpected observation in our study was the divergent tau response after msTBI. Specifically, only a subset of msTBI mice (5/8; 62.5%) showed increased chronic tau burden. The remaining TBI mice (3/8; 37.5%) had a total tau level even lower than that of sham controls. Notably, TBI mice with increased p-tau (PHF1 and AT8) also had increased levels of p62, cathepsin D, and decreased hippocampal size. In contrast, TBI mice with low levels of total tau and tangles also had low p-tau levels and less p62 compared to sham mice. These findings suggest that after TBI, increased tau burden and neurodegeneration may be more associated with the status of the autophagolysosomal pathway, rather than the activity of tau kinase alone.

Macroautophagy (the most well-known form of autophagy) is an essential physiological process that facilitates the breakdown of damaged or toxic cellular components. It involves the formation of double-membrane vesicles known as “autophagosomes” around cargo, which then fuse with lysosomes, forming “autophagolysosomes” and leading to cargo degradation ^60^. One of the major cellular degradation pathways for tau is the autophagolysosome pathway ^61–63^. In humans, many causative genes of frontal temporal dementia (FTD) and amyotrophic lateral sclerosis (ALS) have been identified as important players in the autophagolysosome pathway [Reviewed in ^64^]. In addition, histological examination of brain tissues from patients with tauopathy have demonstrated accumulation of p62-positive, LC3-positive autophagic vesicles and cathepsin D-positive lysosomal vesicles ^40,65^. Neuronal cells derived from induced pluripotent stem cells of FTD patients also showed accumulation of autophagy markers including p62 ^66^. In rTg4510 mice (which overexpress P301L tau), adeno-associated virus (AAV) delivery of transcription factor EB (TFEB, a master regulator of the autophagolysosome pathway) on post-natal day 0 reduced the development of p-tau and NFT and ameliorated neuronal loss ^67^. Mice with a forebrain-specific Atg7 (an essential gene in the macroautophagy pathway) conditional knockout showed age-dependent neurodegeneration and accumulation of ubiquitin-, p62-, and p-tau-positive accumulations ^68^. In Drosophila expressing 2N4R tau, haploid insufficiency of Atg6 (Beclin 1, a critical player of the autophagy pathway) induced neurotoxicity ^69^. In rat hippocampal slices, cathepsin D, a major lysosomal enzyme, degrades tau, and inhibition of lysosomal functions increase tau levels ^70,71^. In addition, in a Drosophila model expressing the R406W mutant tau, genetic deletion of CTSD exacerbated neurotoxicity and shortened lifespan ^72^. These studies demonstrate the central role of the autophagolysosomal pathway in tau clearance and toxicity.

Interestingly, both clinical and animal studies have demonstrated that TBI may disrupt the autophagy-lysosomal pathway and compromise the autophagic degradation activity (also known as “impaired autophagy flux”) ^60^. A clinical case report ^73^ examined 36 autopsy brain samples from patients who died within 1 hr to 7-mo after TBI. They observed increased markers of autophagy, including p62 (SQSTM1) and LC3 cytoplasmic inclusions, within hours to 1-mo post-injury. Another clinical study examined cerebrospinal fluid (CSF) from 30 children (aged 7 weeks to 16 years) at 1d, 3d, and 7d after severe TBI, and compared to age-matched controls ^74^. They found an increase in the autophagy markers beclin1 and p62 in TBI patients. In addition, peak p62 level was associated with unfavorable clinical outcome at 6-mo post-injury, even after adjusting for patient age. In rodent models of TBI and spinal cord injury (SCI), changes in autophagy markers have been reported, including accumulation of beclin1, p62, conversion of LC3-I to LC3-II, and conjugation of ATG12-ATG5 in microsomal fractions ^75–77^ from hours to days post-injury. These studies suggest that impaired autophagy flux is likely initiated in the early phase after neurotrauma, including TBI. In our study, the TBI mice that had increased tau burden also had greater levels of p62 and cathepsin D, suggesting disruption of autophagolysosomal functions in these animals.

The three TBI mice that had low total tau levels also had low levels of markers of autophagolysosomal accumulation, suggestive of increased autophagic flux. Some previous studies have reported reduced neurodegenerative proteins after inflammatory insults, including in one of our previous TBI studies ^78^, where we showed that repetitive mild TBI in 13-mo APP/PS1 mice reduced Thioflavin S+ve amyloid burden at 2-7d post-injury. DiCarlo et al ^79^ showed that intrahippocampal injection of lipopolysaccharide in another APP+PS1 line acutely reduced Aβ immunopositivity at 7d post-injection. Cartagena et al ^80^ showed that penetrating TBI in Sprague Dawley rats reduced full-length tau and increased tau processing and tau fragments at 3d-7d post-TBI. These studies suggest that, in some situations, inflammation may promote turnover of neurodegenerative proteins. The results reported in this study suggests that one possible mechanism leading to increased tau turnover may be through the autophagolysosome pathway.

Our study has several limitations. First, our study design included only one time point. Second, as we did not include mechanistic investigations using inhibitors or enhancers of autophagosomal/lysosomal functions, we cannot conclude whether impaired autophagy-lysosome function is the cause or the result of the divergent post-TBI tau response. There is evidence in the literature supporting both hypotheses. For example, TBI has been reported to impair autophagy flux in many clinical and experimental TBI studies ^60,81^, which could be the cause of tau accumulation. On the other hand, overexpression of tau in mice can lead to impaired autophagy flux ^82^. Further investigation will be needed to determine the cause-and-effect relationship between post-TBI tau burden and autophagy-lysosome function, and understand why this phenomenon only appears in some mice. Third, our study has a small N and may have potential survivor bias in the msTBI group, as we aimed for ~50% mortality (actual 46.7%) and pathological analysis was restricted to surviving mice. Third, it is unclear how the plasma total tau analyses relate to brain tau changes, as high plasma tau levels were observed in mice with normal as well as low tau levels in brain. One possibility is that there is a conformer of brain tau in low tau mice that is not recognized by any of the antibodies used here. Another possibility is that plasma total tau represents both central nervous system (CNS) and non-CNS sources, similar to human studies that report high plasma tau levels after exercise or peripheral injury ^83,84^. Nevertheless, the observation that msTBI appears to accelerate the shift to high plasma tau levels is potentially interesting and will be validated in future studies, particularly with more sampling time points around the point of inflection.

In conclusion, we showed that a single msTBI in rTg4510 mice induced chronic white matter injury, increased GSK-3β activity, and potentially accelerates an increase of plasma tau. Intriguingly, a divergent response in tau species in brain tissue was observed; msTBI induced p-tau in most of the surviving mice, whereas total tau and p-tau levels in others. Interestingly, mice that had increased p-tau also showed accumulation of autophagosomes/lysosomal proteins and decreased hippocampal size, whereas the mice with less total tau showed little autophagolysosome accumulation. The cause-and-effect relationship of post-injury tauopathy and autophagosome/lysosome disruption remains to be elucidated.

## Acknowledgments

We are grateful to the Weston Brain Institute (TR192003) for operating support of this work.

## Figure legends

**Supplementary Figure 1.**
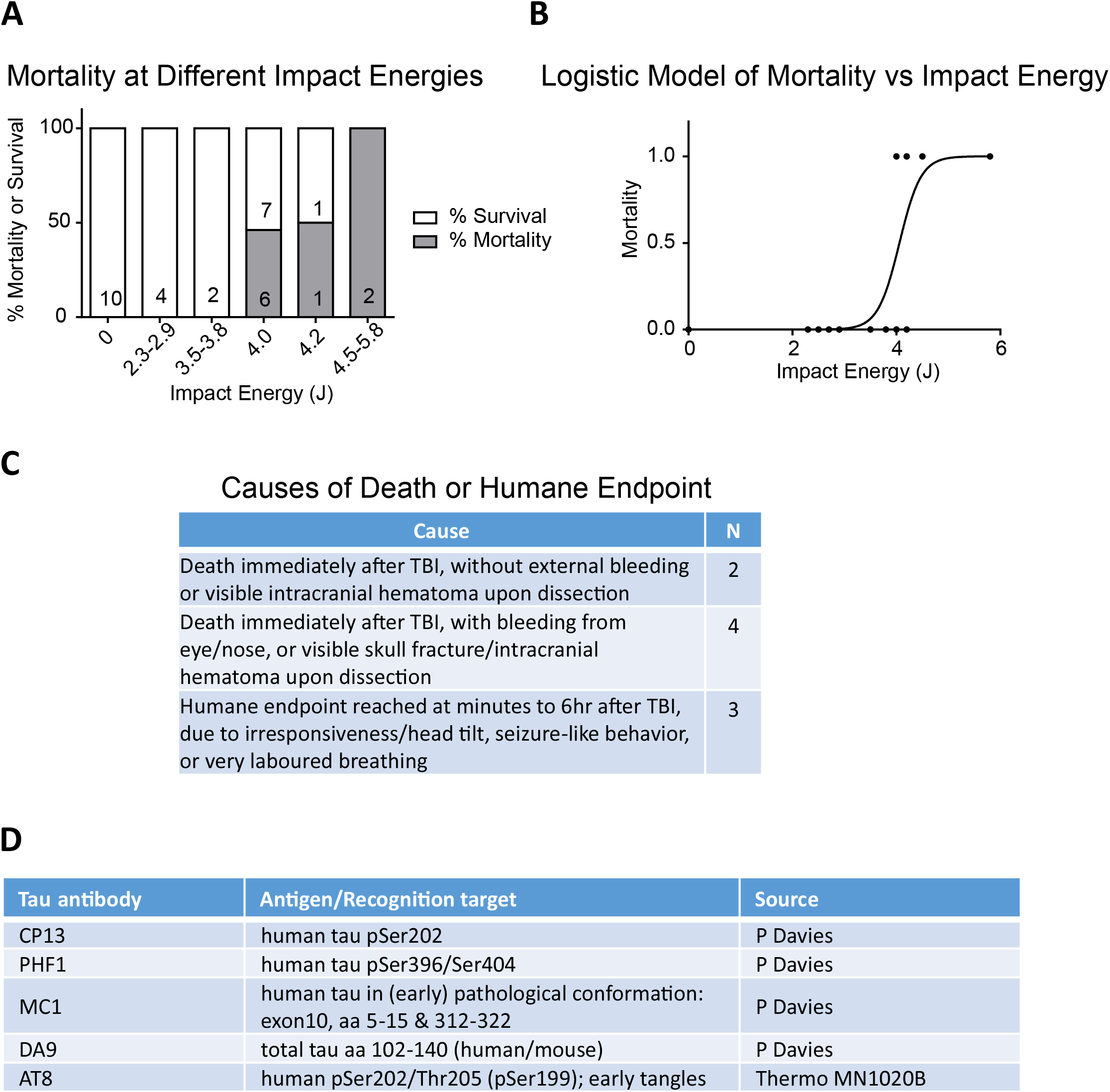
Mortality of 4-mo rTg4510 at different impact energies. (A) Breakdown analysis showing % mortality and % survival at different impact energy ranges. Numbers on the bar represent the number of mice that died/survived from the impact energy range. Note that the 7 mice that survived 4.0J impact and the 1 mouse that survived 4.2J impact were combined into one msTBI group in this study. (B) Logistic regression model of mortality as a function of impact energy. The model is best fitted with the equation y = 1/(1+e^-4.831(x-4.060)^), where y=1 indicates total mortality and y=0 indicates total survival. R^2^=0.4098 (C) Summary of cause of death of animals that did not survive the TBI procedure.

**Supplementary Figure 2.**
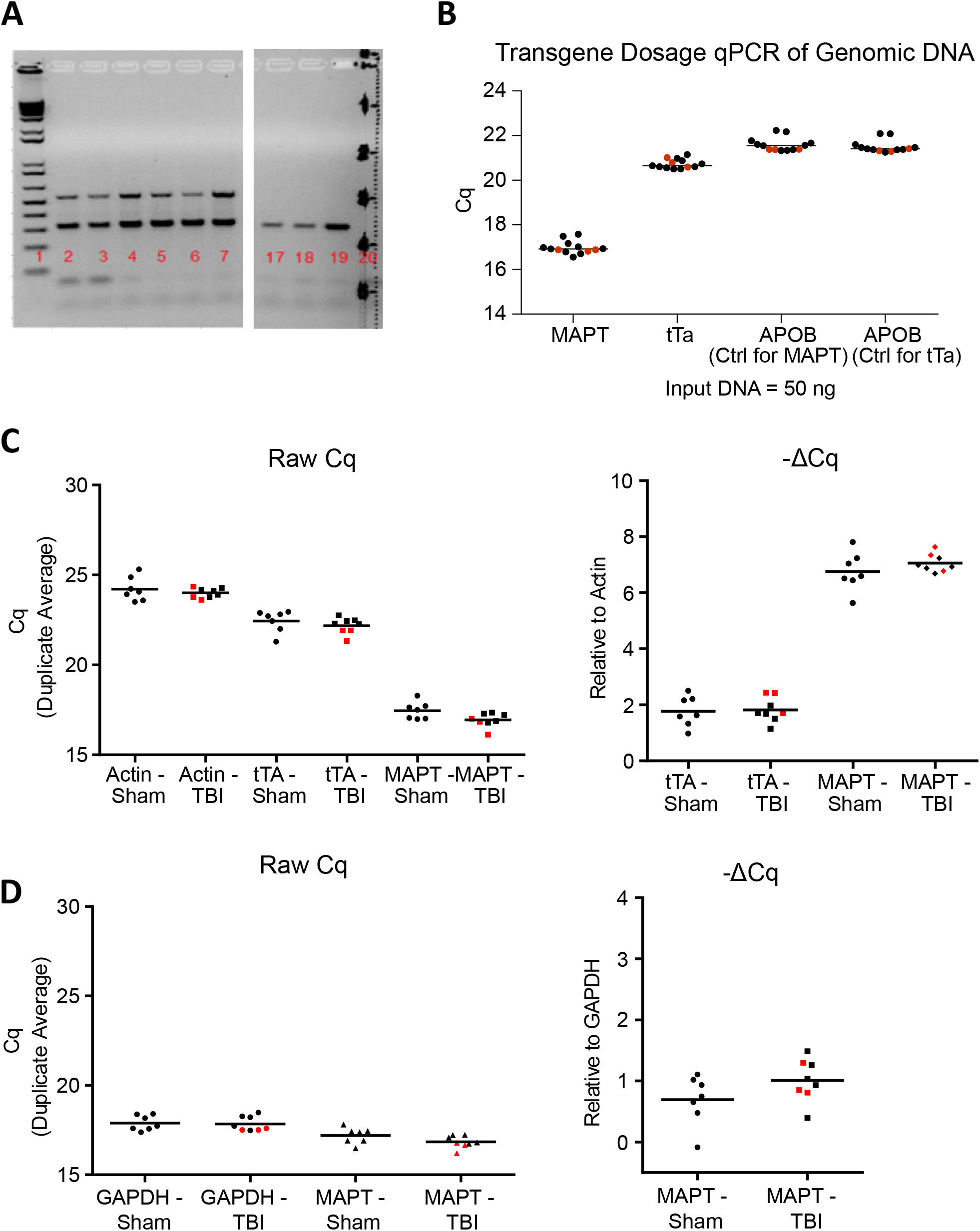
TBI mice with high and low levels of tau did not differ in genotyping, transgene dosage, or transgene expression. (A) Genotyping reconfirmation of rTg4510 mice. Lanes: 1: DNA ladder. 2: Sham rTg4510. 3 and 4: TBI mice with regular level of total tau. 5-7: TBI mice with low tau levels. 17-19: Non-transgenic mice. (B) Transgene dosage was performed using qPCR and confirmed a similar transgene copy number among the entire rTg4510 cohort. (C) Transgene mRNA levels were similar among the entire rTg4510 cohort. (D) Replicate of (C) using a different set of house-keeping gene. Horizontal lines in graphs indicate group mean. Red squares indicate mice with low tau levels.

**Supplementary Figure 3.**
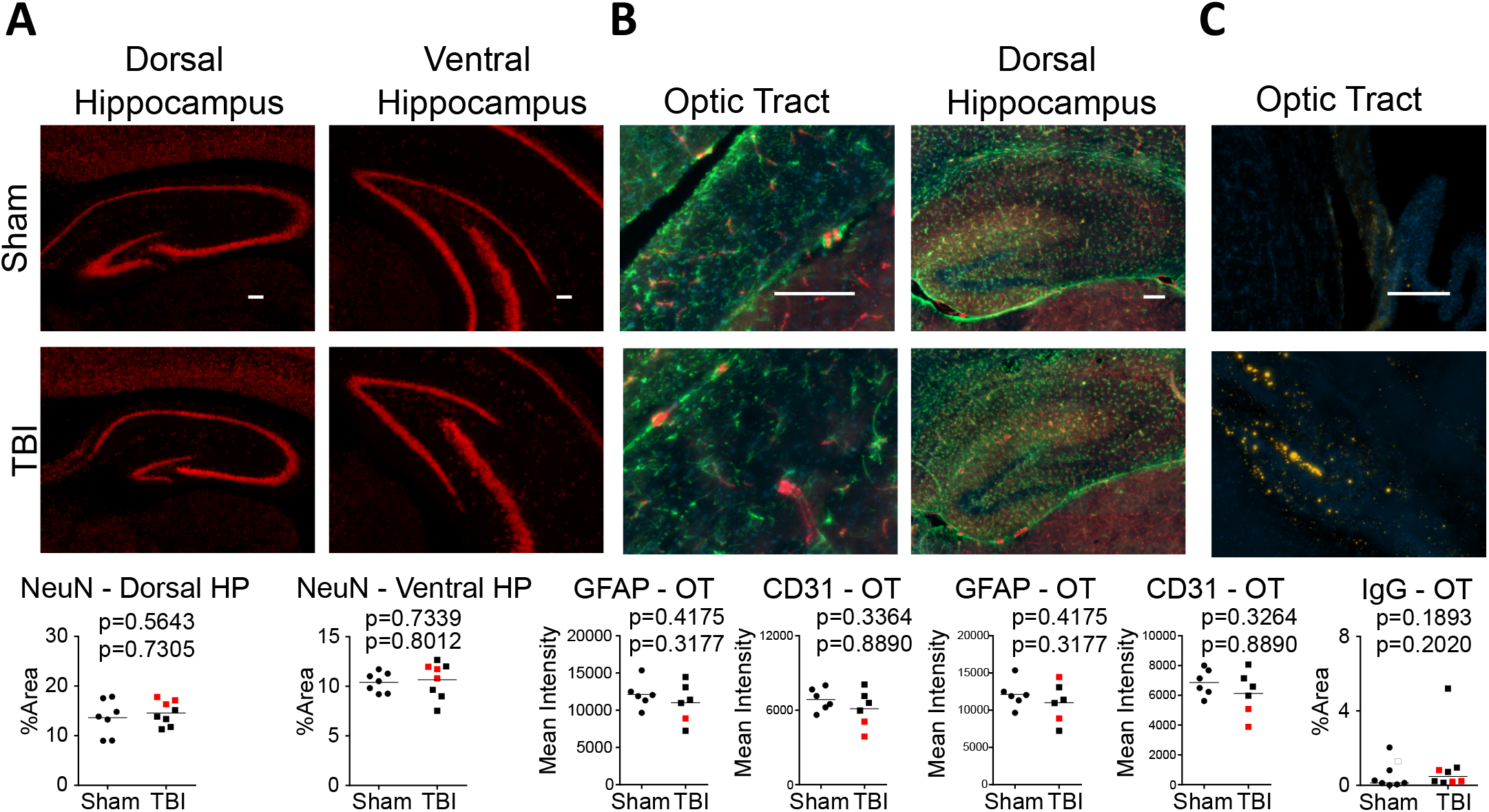
Interfaced TBI does not change levels of neurons, astrocytes, endothelial cells and IgG in rTg4510 mice. Immunohistochemistry of (A) NeuN (red), (B) GFAP (green) and CD31 (red), and (C) IgG (yellow) to stain for neurons, astrocytes, endothelial cells and immunoglobulin G, respectively. DAPI (blue) was stained in (B) and (C) to visualize cell nuclei. Results are quantified in the graphs below the images. Red squares indicate mice with low tau levels. Scale bar = 100 um. Mann-Whitney U test was used for IgG-OT, where horizontal lines in graphs indicate group median. T-tests were used for all other analyses, where horizontal lines in graphs indicate group mean.

**Supplementary Figure 4.**
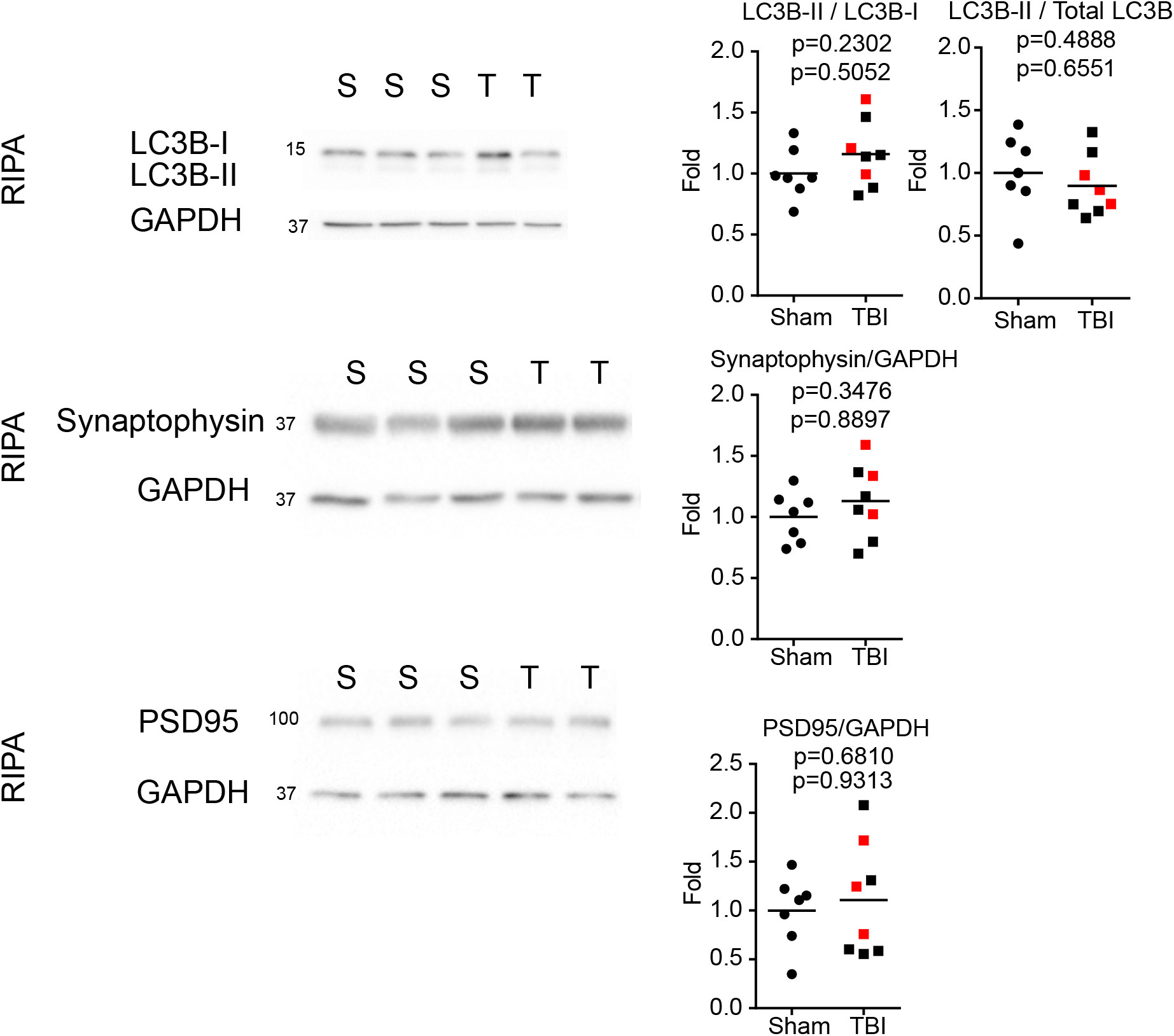
Interfaced TBI does not change protein markers of autophagy initiation or synaptic proteins in rTg4510 mice. Western blotting and quantification of RIPA brain homogenates using antibodies against LC3B, synaptophysin, and PSD95. Red squares indicate mice with low tau levels. T-tests were used for all analyses. Horizontal lines in graphs indicate group mean.

**Supplementary Figure S5.**
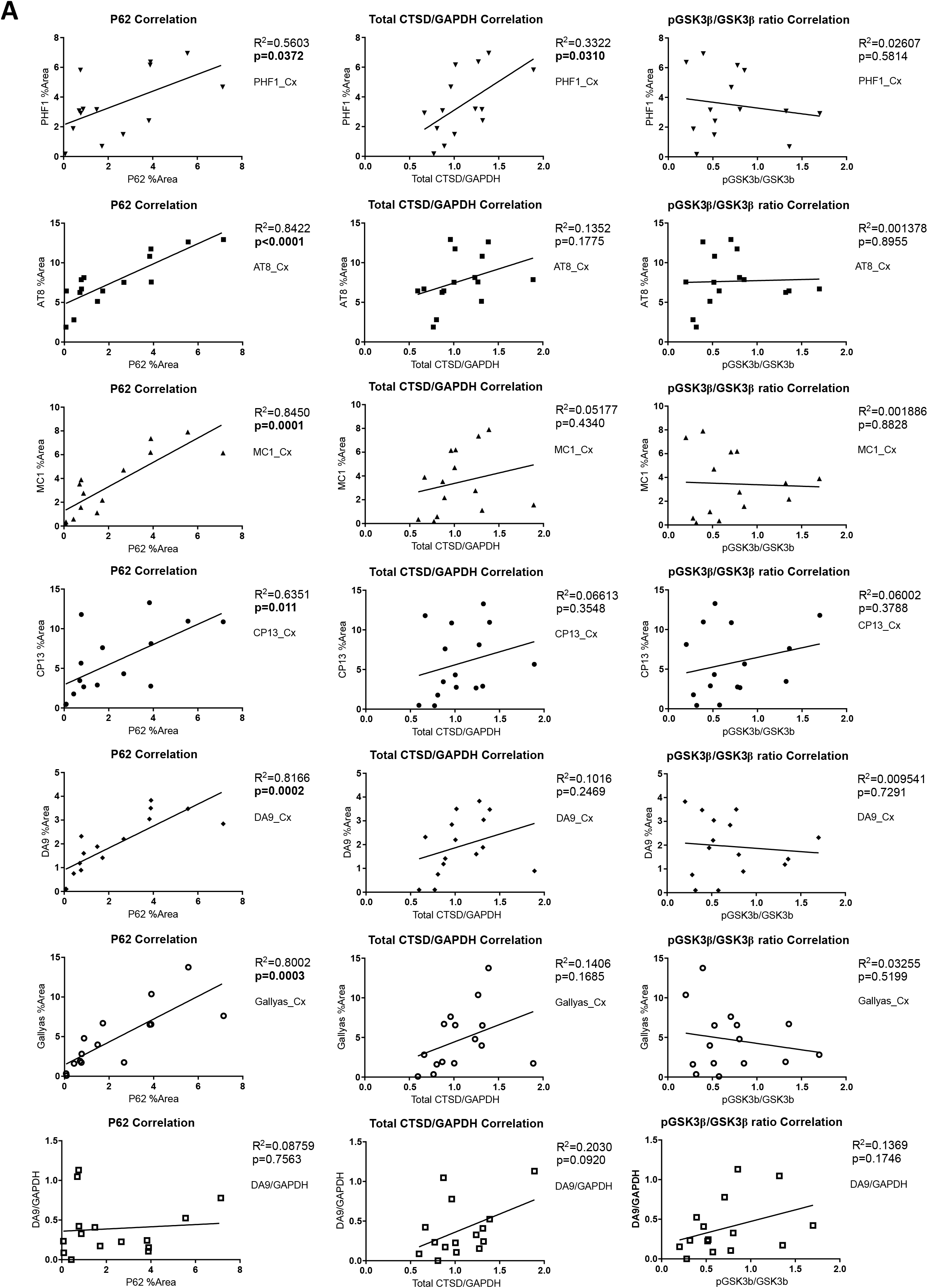
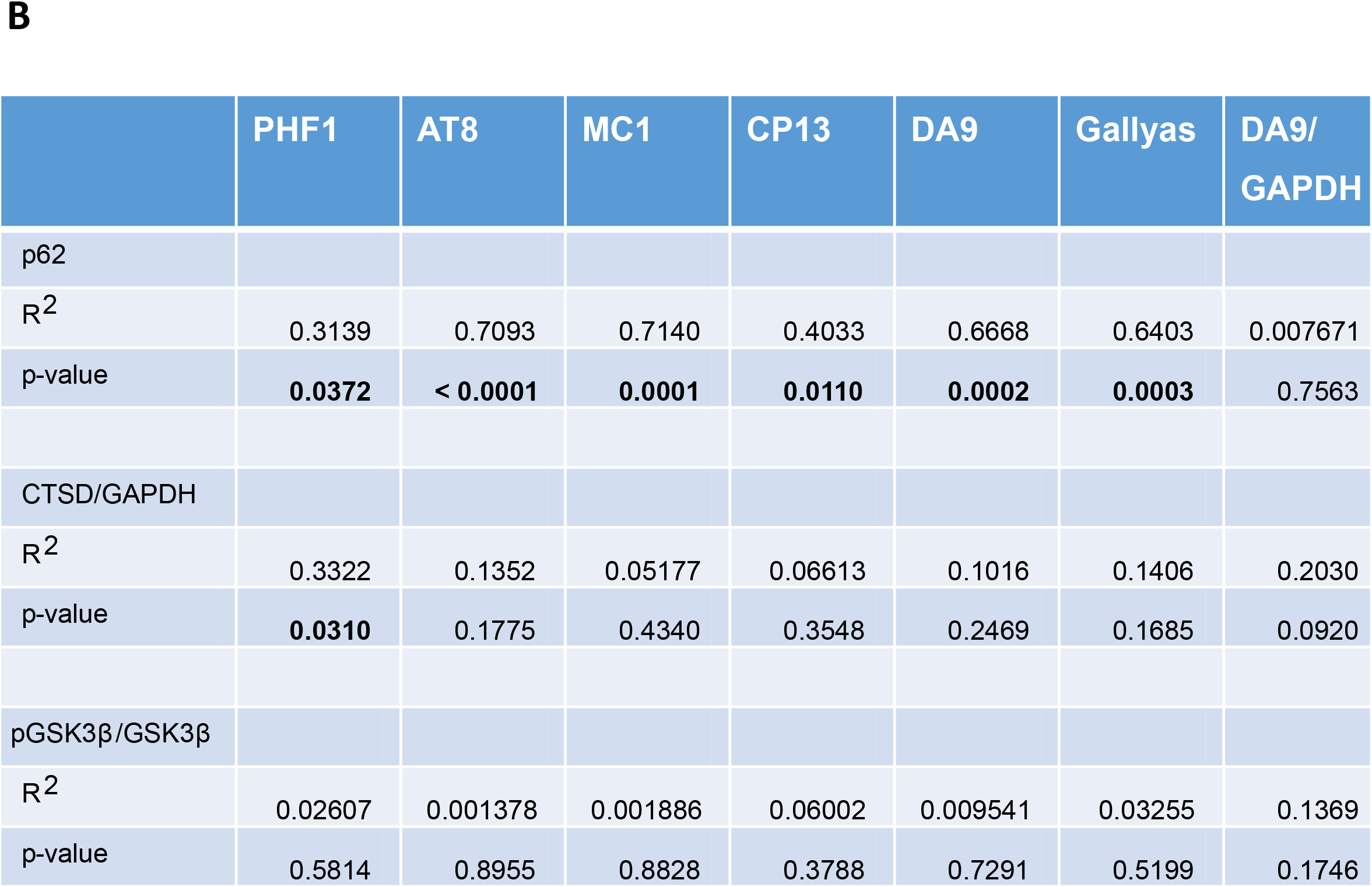
Correlational analyses of tau with autophagolysosomal markers and tau kinase. (A) Plots of Pearson correlation between cortical tau (IHC: PHF1, AT8, CP13, DA9, Gallyas) and tau WB (DA9) with p62, CTSD/GAPDH ratio, and pGSK3b/GSK3b ratio. (B) A summary table of the correlation analyses is shown.

**Supplementary Figure 6.**
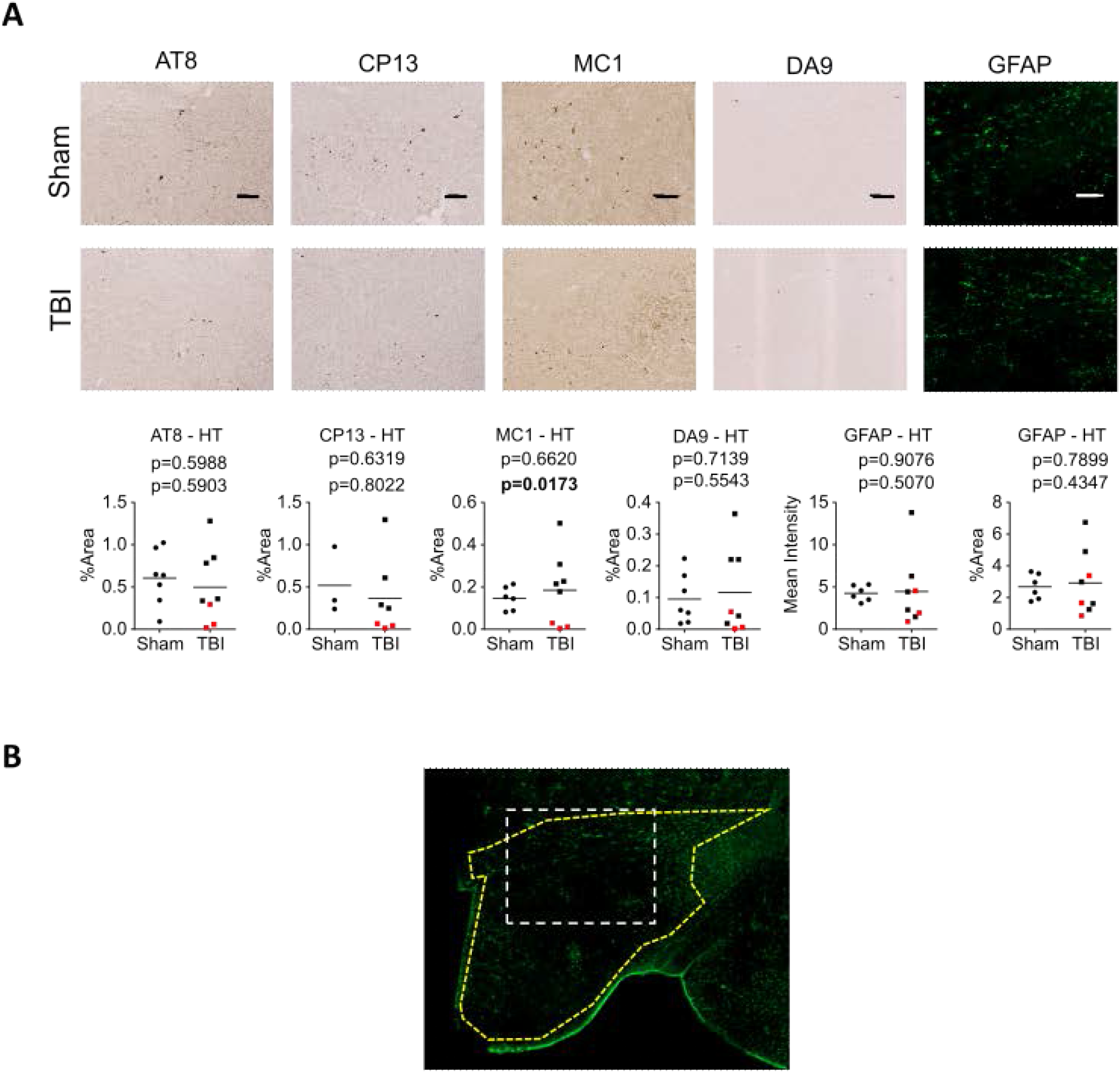
Interfaced TBI did not change tau and astrocyte immunohistochemistry at hypothalamus in rTg4510 mice. (A) Immunohistochemistry of tau (AT8, CP13, MC1, DA9) and astrocyte (GFAP) at hypothalamus were shown. Results are quantified in the graphs below the images. Red squares indicate mice with low tau levels. Scale bar = 100 mm. T-tests were used for all analyses. Horizontal lines in graphs indicate group mean. (B) An example of hypothalamus quantification is shown, using the GFAP-TBI image. The entire hypothalamus region was selected and used for analyses (yellow dotted polygon), and a subset zoomed-in area (white dotted retangle) is shown in (A).

